# Disease networks and their contribution to disease understanding and drug repurposing: Evolution of the concept, techniques and data sources

**DOI:** 10.1101/415257

**Authors:** Eduardo P. García del Valle, Gerardo Lagunes García, Lucía Prieto Santamaría, Massimiliano Zanin, Ernestina Menasalvas Ruiz, Alejandro Rodríguez-González

## Abstract

Over a decade ago, a new discipline called network medicine emerged as an approach to understand human diseases from a network theory point-of-view. Disease networks proved to be an intuitive and powerful way to reveal hidden connections among apparently unconnected biomedical entities such as diseases, physiological processes, signaling pathways, and genes. One of the fields that has benefited most from this improvement is the identification of new opportunities for the use of old drugs, known as drug repurposing. The importance of drug repurposing lies in the high costs and the prolonged time from target selection to regulatory approval of traditional drug development. In this document we analyze the evolution of disease network concept during the last decade and apply a data science pipeline approach to evaluate their functional units. As a result of this analysis, we obtain a list of the most commonly used functional units and the challenges that remain to be solved. This information can be very valuable for the generation of new prediction models based on disease networks.

## 1. Introduction

The study of diseases as non-isolated elements and the understanding of how they resemble and relate to each other are crucial to provide novel insights into pathogenesis and etiology, as well as in the identification of new targets and applications for drugs [1]. The complete sequencing of the human genome at the beginning of the 21st century represented a revolution in the study of the relationships between diseases. In combination with the growing availability of transcriptomic, proteomic, and metabolomic data sources, it should help to improve the classification of diseases [2]. However, the use of these sources raised new problems such as their fragmentation, heterogeneity, availability and different conceptualization of their data [3, 4].

Recent developments in network theory provide a way to address this challenge by representing these complex relationships as a collection of linked nodes [5]. Complex networks theory is a statistical physics interpretation of the old graph theory, aimed at describing and understanding the structures created by the relationships between the elements of a complex system [6–9]. Those elements are represented by nodes, pairwise connected by links whenever a relationship is observed between the corresponding elements. The resulting structure can then be described by means of a plethora of topological metrics [10], or be used as a base for modelling the system. The application of this field to biological problems has been named “network biology”, while its use in biomedical problems is known as “network medicine” [11]. Some applications of biological networks are protein-protein interaction networks, gene regulatory networks (DNA-protein interaction networks), metabolic networks, signaling networks, neuronal network or phylogenetic trees [12].

Following this approach, disease networks express the relationship between diseases as nodes and edges in a graph in *G* = (*D*, *W*), where *D* represents the set of diseases (nodes) and *W* the set of their relationships (edges) based upon their similarity. The meaning of similarity varies depending on the data used to build the network, which may be biological (genes or common proteins) or phenotypic (comorbidity, similar symptoms) [13], among other approaches. As will be explained throughout this article, the concept of disease network is not limited to disease-disease connections (homogeneous networks), but also to relations between the disease and other factors such as its symptoms, its associated genes or its treatments (heterogeneous networks).

During the past decade, numerous studies have been proposed to improve our understanding of the functioning of diseases and their relationships by creating disease networks based on different disease-disease association models and large-scale data exploitation. Of them, a significant number was oriented to exploit the new discovered relationships between diseases in the reassignment of known compounds for their treatment, the so-called “drug repurposing”. In the first part of this document, we will thoroughly review this previous work, analyzing the evolution of the methodologies used in the creation of disease networks from a timeline perspective up to the state of the art.

Despite their different approaches and methodologies, in the studies dedicated to the improvement of the disease understanding and particularly to drug repositioning, the typical phases of a data science pipeline are observed, such as data extraction, data integration model, validation and presentation. In the second part of the document, these common parts are analyzed and their existing implementations are compared. Finally, based on the previous analysis, new studies are proposed by improving or combining the phases of the pipeline.

This work aims to serve as a quick reference guide for practitioners who want to start to create new disease network representations and analyses. Its main contributions are 1) a historical survey of the most relevant studies in this area, 2) a novel comparison of their functional units and 3) a review of the open questions and future lines of research. The study does not intend to evaluate in detail the results of each study or their respective contributions.

## 2. Review methodology

A comprehensive search of the literature on disease networks and their contribution to disease understanding and drug repurposing was undertaken. Five databases were searched, including PubMed/Medline, Web of Science, Springer, ACM Digital Library, and IEEE Xplore. The results were restricted to English-language studies published from 2007 (as described in section 3.1, the earliest study reviewed dates from this year) to September 2018.

The following keywords were used for the search: disease network or at least one of “biological network, network medicine”; and at least one of “disease understanding, drug repurposing, drug repositioning, pipeline”. The obtained studies were screened by a single researcher and then reviewed by the coauthors, so this work can be considered a “systematized review” [14].

Analysis of articles followed predetermined eligibility criteria. Studies were included in the review, if the study: (a) addressed the application of biological networks to disease understanding and/or drug repurposing; (b) d the methods to build the network; (c) provided qualitative and quantitative information of the generated network.

## 3. Evolution of disease networks and their application to drug repurposing

### 3.1. Early studies

Initial works proposing the use of disease networks for the analysis of their underlying relationships exploited data of biological origin. In 2007, Goh et al. constructed a disease-gene bipartite graph called “Diseasome” using information from OMIM database [1]. From the diseasome they derived the Human Disease Network (HDN), in which pairs of disorders are connected if they have common genes. The study revealed that diseases tend to cluster by disease classes and that their degree of distribution follows a power law; that is, only a few diseases connect to a large number of diseases, whereas most diseases have few links to others.

Aiming to reduce the bias of the HDN towards diseases transmitted in a Mendelian manner [15], subsequent studies used other sources of biological data. In 2008 year, Lee et al. constructed a metabolic disease network in which two disorders are connected if the enzymes associated with them catalyze adjacent reactions [16]. In 2009, Barrenas et al. [17] derived a complex disease-gene network (CDN) using GWAs (Genome Wide Association studies). The complex disease network showed that diseases belonging to the same disease class do not always share common disease genes. Complex disease genes are less central than the essential and monogenic disease genes in the human interactome.

The rapid improvement in disease association prediction through the use of network theory fostered its early application to drug repurposing. Drug repurposing is the utilization of known drugs and compounds to treat new indications [18]. Since the repositioned drug has already passed a significant number of toxicity and other tests, its safety is known and the risk of failure for reasons of adverse toxicology are reduced [19]. As a result, the cost and time needed to bring a drug to market is significantly reduced compared to traditional drug development. The commercial applications of drug repositioning and the interest shown by pharmaceutical companies have led to a growing academic activity in this field. This fact is reflected in the evolution of the results for the search by “Drug Repurposing” in Google Scholar, as seen in Figure 1.

**Figure 1.**
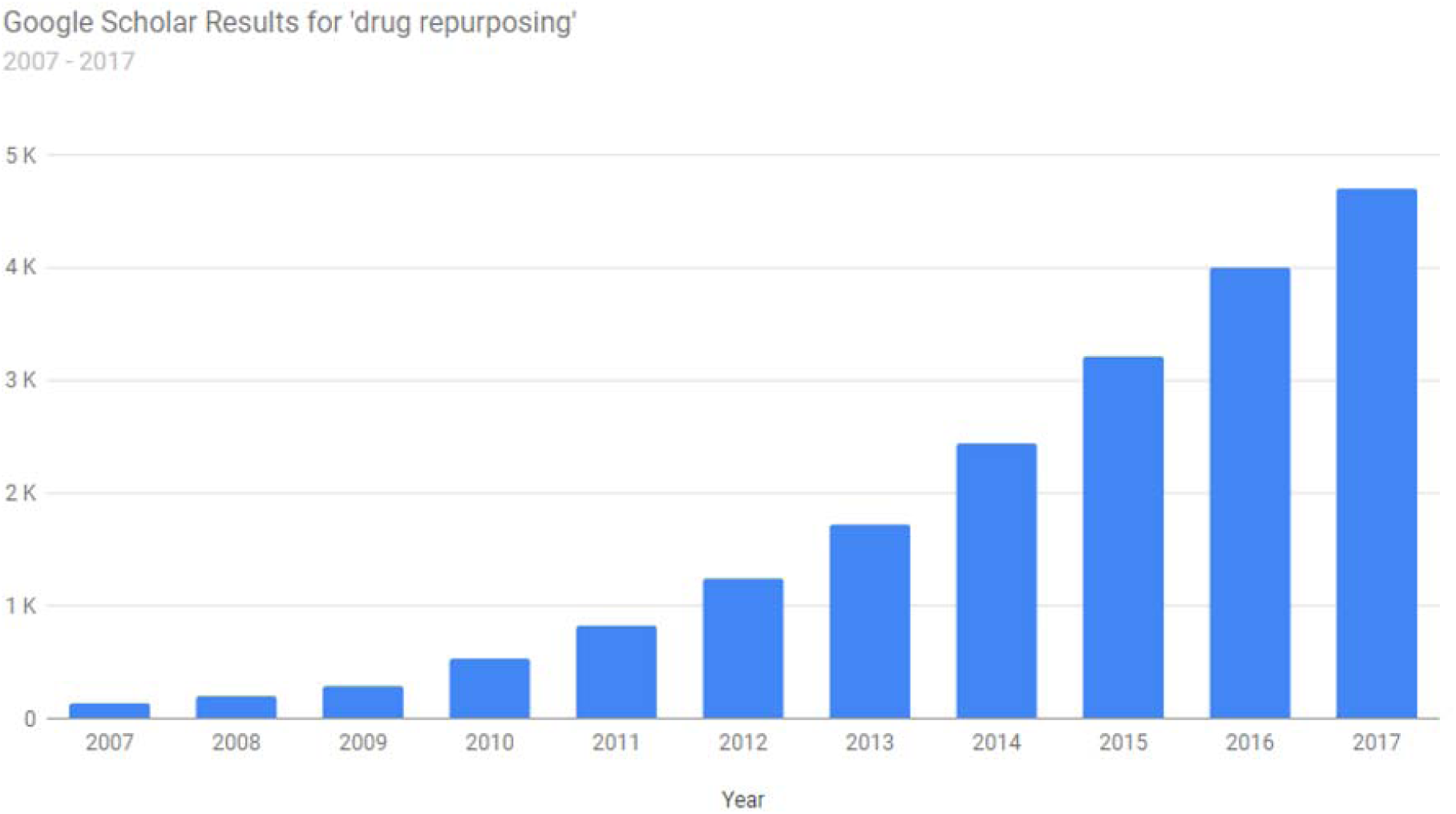
Evolution of the number of articles in Google Scholar containing the term “drug repurposing” within the last 10 years. Retrieved from https://csullender.com/scholar/. Copyright: 2017, Colin Sullender. Use authorized by the copyright owner.

**Figure 2.**
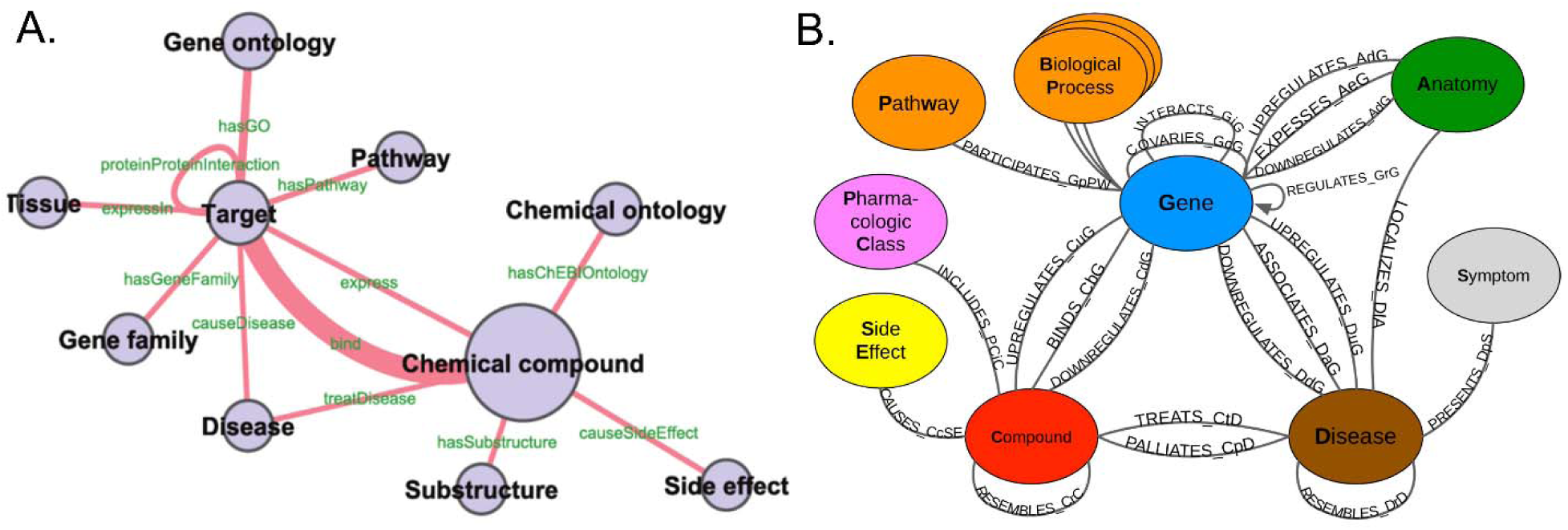
Two examples of semantically annotated heterogeneous networks. Every node and edge was semantically annotated using a systems chemical biology/chemogenomics ontology. In A), Žitnik et al grouped nodes into 10 classes which are linked by 12 types of semantic edges. Two nodes are linked by one or more number of annotated paths. B) shows a Representation of the metagraph in the Hetionet heterogeneous network by Himmelstein et al, with 11 data types (metanodes, depicted as circles) and 24 connection types (metaedges, depicted as links) semantically annotated. Both representations show the capacity of semantic heterogeneous networks to depict the complexity of the relationships between concepts associated with the disease, but they also illustrate the lack of standard in this approach. A. Retrieved from https://journals.plos.org/ploscompbiol/article/figure?id=10.1371/journal.pcbi.1002574.g001. Copyright: 2012 Chen et al. This is an open-access article distributed under the terms of the Creative Commons Attribution License. B. Retrieved from https://www.ncbi.nlm.nih.gov/pmc/articles/PMC5640425/figure/fig1/. Copyright: 2017, Himmelstein et al. This article is distributed under the terms of the Creative Commons Attribution License.

First studies in drug repurposing using biological networks followed a drug-centric approach based on the “guilt-by-association” assumption, that is, similar drugs may share similar targets and vice-versa [20]. In 2007, Yildrim et al. created a graph composed of US Food and Drug Administration–approved drugs and proteins linked by drug–target binary associations [21]. Similar studies were carried out by Ma’ayan [22] in 2007 and Chiang [23] and Bleakley [24] in 2009. In 2008, Nacher Schwartz compiled a drug-therapy network with all US-approved drugs and associated human therapies. Further studies followed a disease-centric approach, in which effective drugs were identified based on disease-disease similarity. In 2008, Campillos et al. predicted new targets for drugs by calculating similarities between diseases based on side effect that appears from injection of drug [25]. In 2009, Guanghui Hu et al. performed a systematic, large-scale analysis of genomic expression profiles of human diseases and drugs to create a disease-drug network [26]. Suthram in 2010 [27] and Mathur in 2012 [28] also predicted new uses of existing drugs based on disease-disease associations calculated from mRNA expression similarity and biological process semantic similarity, respectively.

### 3.2. Incorporating textual sources

The abundance of new biological data did not make researches overlook the existence of another important resource: the highest level clinical phenotypes, that is, symptoms. As one of the first and most obvious forms of diagnosis, the relationship between symptoms and diseases is widely documented in clinical records.

One of the earliest studies in network medicine to use these sources was published by Rzhetsky et al. in 2007. The disease history of 1.5 million patients at the Columbia University Medical Center to infer the comorbidity links between disorders and prove that phenotypes form a highly connected network of strong pairwise correlation [29]. In 2009, Hidalgo et al. built a Phenotypic Disease Network (PDN) summarizing the connections of more than 10 thousand diseases obtained from pairwise comorbidity correlations reconstructed from over 30 million records from Medicare patients. The PDN is blind to the mechanism underlying the observed comorbidity, but it shows that patients tend to develop diseases in the network vicinity of diseases they have already had. Also disease progression was found to be different across genders and ethnicities [30]. More recently, Jiang et al. [31] used data from the Taiwan National Health Insurance Research Database to construct the epidemiological HDN (eHDN), where two diseases are concluded as connected if their probability of co-occurring in clinics deviates from what expected under independence.

Despite their demonstrated potential in pathological analysis, the access and use of clinical records in medical research is limited by several issues, including the heterogeneity of sources [32], ethical and legal restrictions and the disparity of regulations between countries [33]. The analysis of open text sources has been used as an alternative to medical records. One of the reasons is the improvement in the techniques for Named Entity Recognition (NER) for the extraction of medical terms.

Okumura et al. [34] performed an analysis of the mapping between clinical vocabularies and findings in medical literature using OMIM as a knowledge source and MetaMap as the NLP tool. Following this idea, Rodríguez et al. [35] used web scraping and a combination of NLP techniques to extract diagnostic clinical findings from MedlinePlus articles about infectious diseases using MetaMap tool. In a further study, the same team compared the performance of MetaMap and cTakes in the same task [36].

The increasing availability of retrieval engines such as PubMed or UKPMC, maintained by the US National Center for Biotechnology Information (NCBI) and the European Bioinformatics Institute (EBI), respectively, has also boosted this approach [37]. In 2014 Zhou et al. extracted symptom information from PubMed to construct the Human Symptoms Disease Network (HSDN). In the HSDN, the link weight between two diseases quantifies the similarity of their respective symptoms [38]. In 2015, Hoehndorf et al. created yet another Human Disease Network using a proposed similarity measure for text-mined phenotypes [39]. In both cases, these studies compare their results with gene-based networks, finding that symptom-based similarity of two diseases strongly correlates with the number of shared genetic associations. They also demonstrated that not only Mendelian diseases tend to be grouped into classes, but also common ones.

### 3.3. Completing the diseasome

Due to the intrinsic complexity of the relationships between diseases, the consideration of a single factor (e.g. gene-disease, symptom-disease or drug-disease) was a limiting factor to obtain novel findings and predicting drug repositioning [18]. In his review of the HDN in 2012, Goh. et al. proposed that each and every disease-contributing factor such as molecular links from interactome, co-expression and metabolism, as well as genetic interactions and phenotypic comorbidity links, will have to be integrated in a context-dependent manner. Furthermore, drug chemical information and non-biological environmental factors such as toxicity information altogether must also be incorporated [15]. The result will be a combination of general and bipartite network representations into a single, complex, k-partite heterogeneous network referred as the complete Diseasome.In line with this idea, Gottlieb made use of a broader collection of data sources to create five drug-drug similarity measures and two disease-disease similarity measures. These similarity measures were then used by PREDICT, an algorithm to infer novel drug indications [40]. Further studies by Sun [41], Albornoz [42], Daminelli [43] and Wang [44] combined multiple data sources to create tripartite networks of gene-disease-PPI, gene-disease-pathways and drug-target-disease, to predict disease-disease associations and repurposing candidate drugs. In 2012, Chen et al. created an heterogeneous network from 17 public data sources relating to drugs, chemical compounds, protein targets, diseases, side effects and pathways [45]. In 2013, Žitnik et al. integrated molecular interaction and ontology data of 11 different types to create another heterogeneous network. When evaluating the predictive capacity of the network, genetic interactions proved to be the most informative feature, as they tend to be causative as opposed to correlative and may therefore have less noise associated [4]. In both studies, the authors leveraged semantic ontology-level information to annotate the edges. The evolution of these type of heterogeneous networks has resulted in the generation of complex tools for the study of disease associations based on multiple sources and types of relationships. A notable example is Hetionet [46], an integrative network encoding knowledge from millions of biomedical studies. Its data were integrated from 29 public resources to connect compounds, diseases, genes, anatomies, pathways, biological processes, molecular functions, cellular components, pharmacologic classes, side effects, and symptoms. However, the main disadvantage of this approach is the lack of standards and the ambiguity in node/link descriptions, as shown in **Figure 1**, which may lead to wrong inferences.

Ultimately, advances towards more comprehensive networks have resulted in tools for the prediction of new treatments given a certain disease, known as Drug Target Indications (DTI). This is the case of Rephetio [47], a project based on Hetionet [46] that predicted 3394 repurposing candidates by applying an algorithm originally developed for social network analysis [48]. Another example worth mentioning is DTINet [49], a DTI prediction system based on learning low-dimensional feature vectors that showed better performance than other state-of-the-art prediction methods and discovered the potential application of cyclooxygenase inhibitors in preventing inflammatory diseases.

## 4. A data science pipeline to build disease networks

Throughout the previous section, we have seen how the rise of network medicine studies has resulted in a expanding variety of innovative methods for the construction and exploitation of disease networks. However, despite using different strategies, these methods are generally based on determining the similarities and relationships between diseases and their treatments at phenotypic level (comorbidity, side-effects) or biological level (common genes, proteins, compounds). Furthermore, they clearly share common phases such as data ingestion, data processing, analysis, modeling or visualization that can be represented as functional units of a data science pipeline.

The data science pipeline consists in a sequence of stages or functional units that sequentially process some input data in order to solve a certain problem [50]. This concept applies to disease networks, where disease information is processed to discover how diseases relate to each other or how drugs can be repositioned. The pipeline representation also facilitates the reproducibility and the comparison among studies as a whole and also at phase level. Most importantly, it also enhances the reusability and the recombination of the functional units to build new drug-repurposing. Throughout the following subsections, we will describe the process of construction and exploitation of a disease network through the functional units of a data science pipeline.

### 4.1. Data acquisition and processing

The first step in the pipeline is to acquire data from a variety of sources, a process known as data acquisition or data ingestion. As seen in the section 2, the growing availability of information sources has allowed developing different approaches to improve our understanding of diseases and to predict new drug applications.

A significant number of studies use biological data, such as KEGG (genes and pathways) [16], BioGRID (protein interactions) [4] or OMIM (genes and phenotypes) [1, 28], among many others. **Supplementary Table 1** contains some of the most important sources of biological information, including their type and description. Studies on disease networks focusing on drug repositioning exploit drug databases and their relation to genes, phenotypes and compounds, such as those offered by the FDA [21–24] or DrugBank [25–28, 38], for instance. **Supplementary Table 2** collects the most common drug data sources. Finally, an increasingly significant number of studies use data obtained by mining medical literature sources (e.g. articles, clinical trials) such as PubMed [38, 39, 51] or the GWAS Catalog [41]. **Supplementary Table 3** contains some of the most relevant sources of medical literature.

A second step in the pipeline consists in transforming and mapping data into a format with the intent of making it more appropriate to work (usually referred as data processing, data wrangling or data munging). Recent studies combine multiple databases to provide more accurate prediction models [4, 45, 46]. However, this poses a challenge when relating identifiers or terms obtained from different sources. To address this problem, researchers use thesauri of terms such as MeSH, SNOMED CT or UMLS; code listings such as ICD or HGNC; and ontologies such as DO, PO, GO or Uberon [39, 52]. Being a valuable source of semantic and hierarchical information themselves, these resources allow mapping data such as disease codes or medical terms. In the case of medical literature sources, the use of metadata (such as MeSH headers in the case of Pubmed, for example) is often combined with terms extraction tools such as MetaMap or cTakes [36]. **Supplementary Table 4** lists some of the sources used for data mapping.

The way to exploit the information in these databases varies greatly from one source to another. Largest databases offer online advanced search and provide developers with application programming interfaces (APIs) to facilitate intensive access to data. For example, the NCBI provides the E-utilities, a public API to access all the Entrez databases including PubMed, PMC, Gene, Nuccore and Protein. The Japanese KEGG also provides REST APIs for data consumption. DisGeNET provides an SPARQL endpoint that allows exploration of the DisGeNET-RDF data set and query federation to expand gene-disease association information with data on gene expression, drug activity and biological pathways, among other. In some cases, data can also be downloaded for their consumption through on-premise applications, as in the case of the Disease Ontology or the Gene Ontology, for example. This disparity complicates the use of different sources in research projects. To alleviate this problem, initiatives such as Biopython^1^ offer common libraries to access multiple sources reducing code duplication in computational biology. Finally, it is very important to know the limitations imposed by each source regarding the volume and use of the data. **Supplementary Tables 1-4** also include information in this regard.

### 4.2. Data integration and modeling

In the next steps of the data science pipeline, data previously acquired and processed are integrated and analyzed in order to answer the matter of our study. In other words, a disease network is built by combining the output of the previous stage and a model is constructed from it. Disease networks consist of a set of nodes (mainly, but not only, representing diseases) and a set of edges (connecting diseases directly or through other related node types). Depending on the type of node they connect, network edges can be directed or undirected, weighted or unweighted. As described in previous sections, over the past decade successive studies based on disease networks have proposed different models of data integration.

#### 4.2.1. Homogeneous networks

Homogeneous disease networks (i.e. those where nodes represent diseases and edges represent direct connections among them) are the simplest type of disease networks. In many studies these networks are built as a projection of a heterogeneous disease network (i.e. a network in which diseases are connected to other types of nodes) [1, 42]. For example, in **Figure 3**, the gene-disease bipartite network is projected onto the disease similarity network (DSN) by relating two diseases that have a gene in common. The disease–disease network can then be analysed by using standard network based methods [1, 53]. In a simplistic approach, the link weights in the resulting disease–disease network represent the link multiplicity resulting from the projection. More complex methods, such as hyperbolic weighting or resource allocation weighting, have been proposed as an alternative [54, 55].

**Figure 3.**
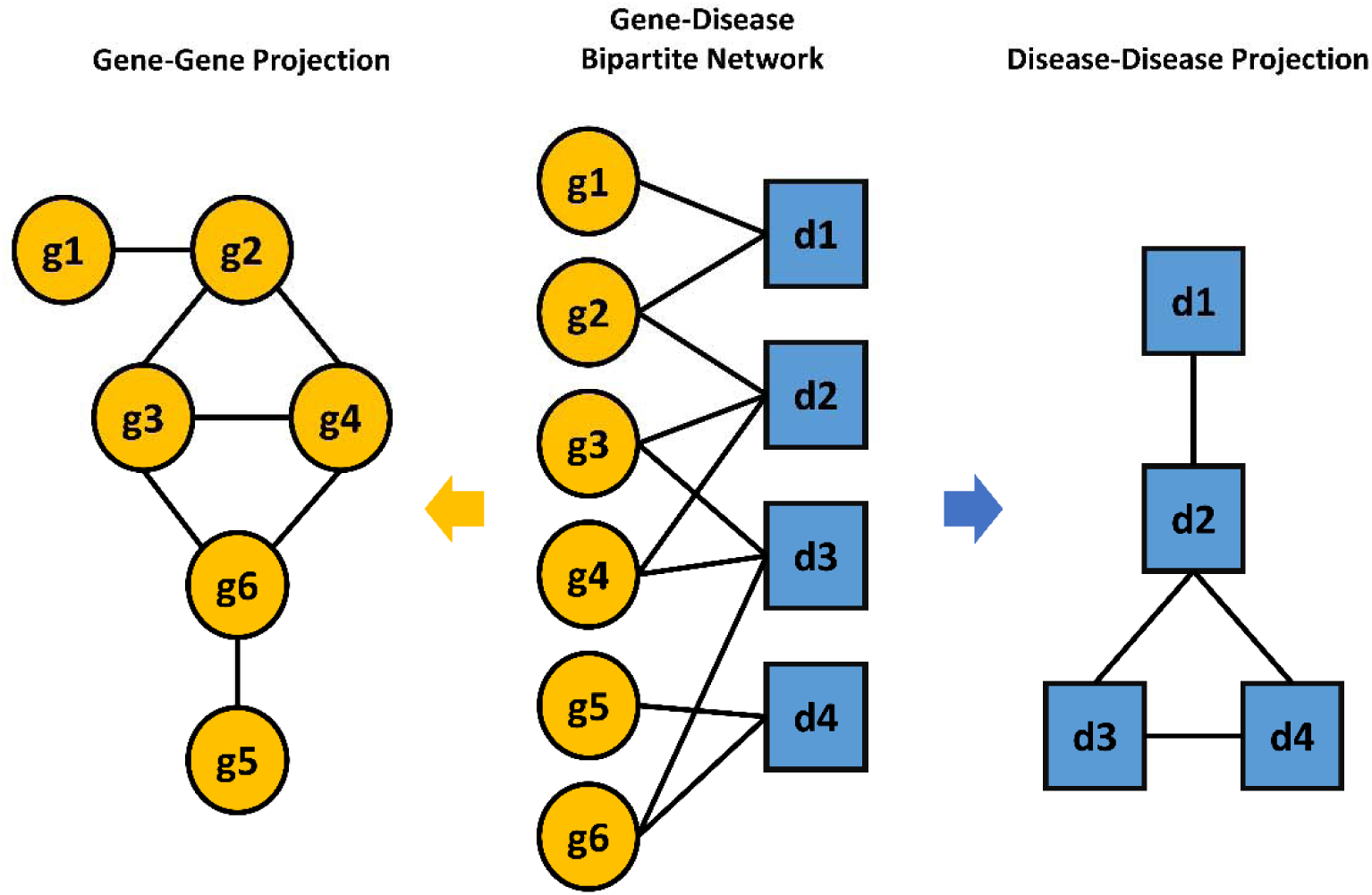
A representation of a heterogeneous network composed of gene-disease interactions. Homogeneous gene-gene or disease-disease networks are obtained via gene or disease projections, respectively.

In other studies, homogeneous disease networks are built as similarity networks. In these networks, if the similarity score between disease i and j is more than zero, the corresponding vertices are linked by an edge in the network. The weight of this edge is the corresponding disease similarity score. Several computation methods for the disease similarity score have been proposed, being Vector Space Model (VSM) [56] among the most popular ones. For instance, in 2006 Van Driel et al. represented diseases as vectors of features (viz. disease associated MeSH terms extracted from OMIM records) weighted by their inverse document frequency [57]. The similarity between diseases was then computed as the cosine of the disease vector angles (i.e. cosine similarity). A similar approach was followed by Zhou to build the HSDN [38] and by Sun to build the Integrated Disease Network [58]. Hoehndorf et al. proposed Normalized Pointwise Mutual Information (NPMI) for disease phenotypic term weighting and later used the PhenomeNET system to compute similarity between diseases using a Jaccard index based measured [39]. Similarity measures based on the term hierarchy in the Disease Ontology and the Gene Ontology have been proposed by Resnik, Lin, Wang, Mathur and Cheng [28, 59–62], and have been integrated in online tools like DisSim or DisSetSim [63, 64]. Okumura et al. described alternative similarity measures based on standardized disease classification, probabilistic calculation, and machine learning [65].

#### 4.2.2. Heterogeneous networks

The projection of heterogeneous networks into homogeneous disease-disease networks allows applying simpler network analysis techniques on the resulting network. However, it often results in information loss. For instance, in **Figure 3** by projecting the gene-gene network onto the disease-disease network, the information about gene interactions and their structure is lost. In contrast, heterogeneous networks make it easy to predict relationship between entities of different types, such as diseases, genes or drugs, following a guilt-by-association paradigm [20]. For example, a drug that regulates a gene associated to a disease could be repurposed for diseases associated to the same gene. Data fusion by matrix factorization and network topology based techniques, such as diffusion and meta-path, are the most common methods for edge prediction in heterogeneous networks, as discussed below

Matrix Factorization methods are closely related to clustering (unsupervised) algorithms. Non-Negative Matrix Factorization (NNMF) decompose matrices of heterogeneous data and data relationships to obtain low-dimensional matrix factors. These factors are then used to reconstruct the data matrices, adding new unobserved data obtained from the latent structure captured by the low-dimensional matrix factors. Hence, NNMF provides a mechanism to integrate heterogeneous data of any number, type and size. In 2013 Žitnik et al. applied a variant of NNMF called non-negative matrix tri-factorization to discover new disease-disease association by fusing 11 data sources on four type of objects including drugs, genes, DO terms and GO terms [4]. In 2015 Dai et al. integrated drug-disease associations, drug-gene interactions, and disease-gene interactions with a a matrix factorization model to predict novel drug indications [66]. More recently, Zhang et al. proposed a similarity constrained matrix factorization method for the drug-disease association prediction using data of known drug-disease associations, drug features and disease semantic information [67].

Methods based on diffusion (i.e. information spreading across network links) have also been extensively proposed to estimate the strength of the connection between nodes of heterogeneous networks. An advantage of such approaches, also called network propagation methods, over matrix factorization is that they they preserve the network structure. Chen et al developed the method of Network-based Random Walk with Restart on the Heterogeneous network (NRWRH), a variation of a ranking algorithm, to predict potential drug-target interactions on heterogeneous networks [68]. Further variations of random walk algorithms, such Bi-Random Walk (BiRW) have been applied to predict novel disease-gene [69], disease-MiRNA [70] or disease-lncRNA associations [71], among others.

Metapath-based approaches also preserve the network structure, and additionally provide an intuitive framework and interpretable models and results. A meta-path P is a path defined over the general schema of the heterogeneous network G = (A, R), where A represents the set of nodes and R the set of their relationships. The metapath is denoted by 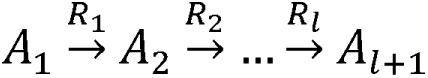, where *l* is an index indicating the corresponding metapath [48]. **Figure 4** shows the metapaths extracted from an annotated heterogeneous network.

**Figure 4.**
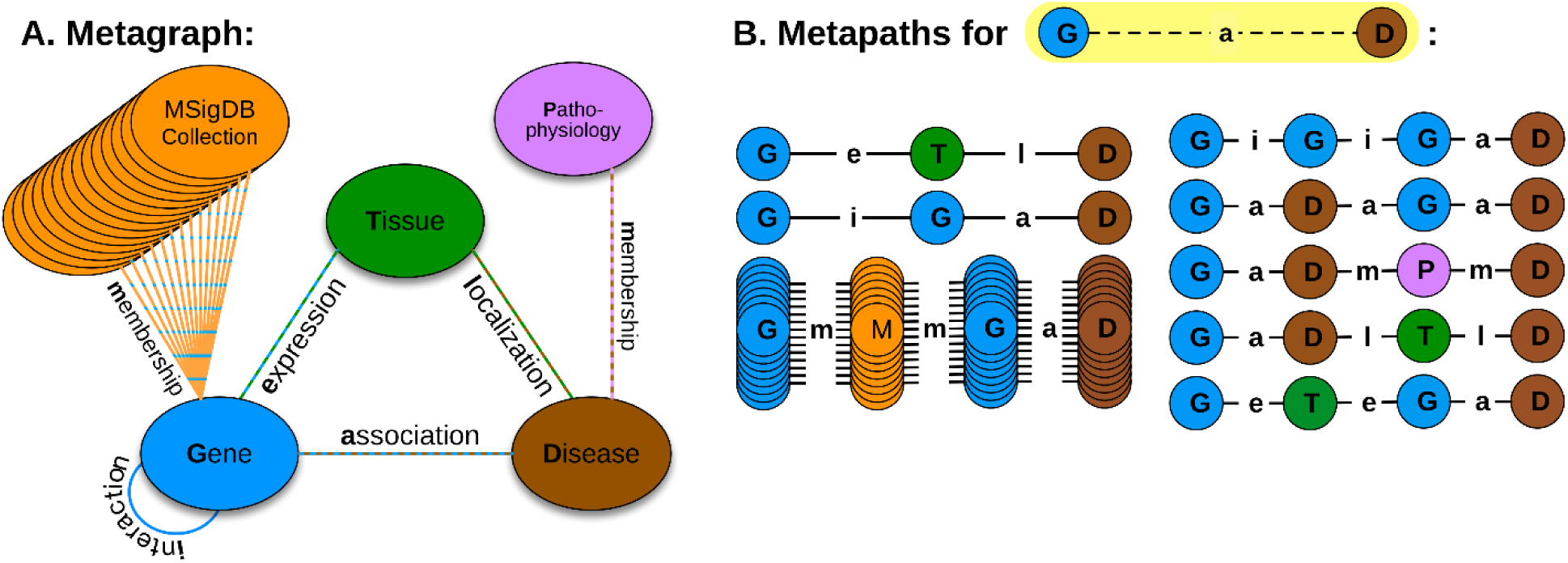
A) The Hetionet annotated heterogeneous network is constructed according to the metagraph schema, which is composed of metanodes (node types) and metaedges (edge types). B) The network topology connecting a gene and disease node is measured along metapaths (types of paths). Starting on Gene and ending on Disease, all metapaths length three or less are computed by traversing the metagraph. Retrieved from https://journals.plos.org/ploscompbiol/article?id=10.1371/journal.pcbi.1004259. Copyright: 2015 Himmelstein, Baranzini. This is an open access article distributed under the terms of the Creative Commons Attribution License.

In their 2012 study, Chen et al. developed a meta-path based statistical model called Semantic Link Association Prediction (SLAP) to assess the association of drug target pairs and to predict missing links [45]. In 2016 Gang Fu et al. proposed an alternative DTI approach to the SLAP algorithm taking advantage of machine learning methods such as Random Forest and Support Vector Machine [64]. To quantify the prevalence of the meta-paths, Himmelstein adapted an existing method developed for social network analysis (PathPredict) and developed a new metric called degree-weighted path count (DWPC). The DWPC downweights paths through high-degree nodes when computing meta-path prevalence [46].

Despite maintaining and exploiting the structure of heterogeneous networks, methods based on diffusion or meta-paths present some scalability limitations, such as the bias introduced by the noise and high-dimensionality of biological data or the effort in feature engineering. Recently, Luo et al. designed DTINet, a novel network integration pipeline for DTI prediction. DTINet integrates information from heterogeneous sources (e.g., drugs, proteins, diseases and side-effects) and copes with the noisy, incomplete and high-dimensional nature of large-scale biological data by learning low-dimensional but informative vector representations of features for both drugs and proteins [49].

### 4.3. Model validation

In this analysis of the reconstruction of disease networks, we wanted to give a special relevance to the validation process. Ensuring that the computational pipeline is producing correct and valid results is critical, particularly in a clinical setting [72]. As previously explained, disease networks are used in studies as diverse as the discovery of new disease-disease relationships, the prediction of gene-disease relationships (GDA) or the repositioning of drugs. The validation of the network depends, therefore, on the type of study in question. In general, the validation can be done experimentally or by computational techniques.

#### 4.3.1. Approaches

Experimental validation includes the verification of the predictions in a controlled environment outside of a living organism (*in vitro*) or using a living organism (*in vivo*). Animal studies and clinical trials are two forms of in vivo research. For example, in their drug repositioning study based on heterogeneous networks, Luo et al. validated the bioactivities of the COX inhibitors predicted by DTINet experimentally. They tested their inhibitory potencies on the mouse kidney lysates using the COX fluorescent activity assays [49]. Jodeleit et al. validated their disease network of inflammatory processes in humanized NOD/SCID/IL2R_γ_ (NSG) mices [73]. While experimental validation studies have the potential to offer more conclusive results about the performance of disease networks, they have several limitations. First, animal studies and clinical trials require expensive lab work and are long and costly. In addition, their conclusions can be misleading. For example, a therapy can offer a short-term benefit, but a long-term harm. Finally, the relevance of animals as models of human disease is questionable, because the networks linking genes to disease are likely to differ between the two species [74, 75].

*In silico* is an expression used to mean “performed on computer or via computer simulation.” In silico tests have the potential to speed the validation process while reducing the need for expensive lab work. In silico validation requires a point of reference for evaluating the model performance, also known as *Criterion Standard* or *Gold Standard*. It is noteworthy that in the field of biomedicine usually the *criterion standard* is actually the best performing test available under reasonable conditions [76]. For example, in this sense, a MRI is the gold standard for brain tumour diagnosis, though it is not as good as a biopsy [77]. Hence, the most recurrent benchmarks used in the validation in silico of disease networks include consolidated data biomedical sources and medical literature.

While in silico validation is especially appealing because of the ability to rapidly screen candidates and to reduce the number of possible repositioning candidates, it is debatable how useful and reliable such methods are in producing clinically efficacious repositioning hypotheses. With in silico repurposing approach, there can be a possibility of false positive hits during screening and also the activity of the candidate drug molecules may vary in the in vitro or in vivo systems. Therefore, to validate their potency further in vitro and in vivo studies are needed to be performed [78].

#### 4.3.2. Sources

Sources of biological, phenotypic or chemical data as well as several available ontologies and code standards (see section 3.1) are used for in silico validation in many studies focusing on disease networks. For instance, their performance to discover disease-disease relationships has been validated with the disease classifications in the Disease Ontology [4, 39] or in the ICD codes [42], as well as with comorbidity associations downloaded from the Human Disease Network (HuDiNe) [41]. DisGeNET has been used to validate de novo gene-disease associations [79], as it integrates data from expert curated repositories with information gathered through text-mining of the scientific literature, GWAS catalogues and animal models [80].

For the validation of drug repositioning predictions, sources such as PharmacotherapyDB and DrugCentral were exploited [47]. However, the heterogeneity in the source of these standards as well as the types of data they contain is detrimental to reproducibility and may lead to claims on extremely high accuracy. For instance, DrugBank contains information about only the FDA-approved indications for drugs, and missesit misses off-label uses and late-stage clinical trials. On the other hand, Comparative Toxicogenomics Database contains literature-annotated links between drugs and both approved and investigational indications, and contains drug–indication pairs that have subsequently failed to receive FDA approval [81].

Furthermore, the aforementioned sources are inevitably biased towards consolidated knowledge, and therefore they might suffer some limitations in corroborating new discoveries. As an alternative (or usually, as a complement) to these sources, medical literature (i.e. studies, medical trials, clinical histories) are used for *in silico* validation of disease network based studies. For instance, Mathur and Paik used previous studies to validate disease-disease and drug-target associations [28, 82]. In some cases, the validation process also combined expert review to corroborate the discoveries [38].

#### 4.3.3. Methods

Leaving aside the particularities of biomedical research and its sources, the validation of classification or prediction methods based on disease networks does not differ from other validation cases. Therefore, in the analyzed studies we found validation methods widely used. For example, k-fold cross-validation is often used to check whether the model is an overfit or not [83, 84]. Overfitting is one of the typical problems of validation, especially when limited data sets are available. Information leakage is another common pitfall in model validation, and occurs when biased data (e.g. data on the training labels) leak into the model before cross-validation, making irrelvant features appear as highly predictive and leading to overly optimistic results. This issue can be prevented by applying nested cross-validation or by excluding certain datasets before the validation process [85, 86].

To quantify the predictive power of their network-based model, many studies use the Area Under the Curve of the Receiver Operating Characteristic (AUC-ROC), another frequently used method in validation problems [39, 87, 88]. The AUC-ROC is the plot between sensitivity and (1-specificity). The *p*-value is the probability that the observed sample AUC-ROC could actually correspond to a model of no predictive power (null hypothesis), i.e. to a model whose population AUC-ROC is 0.5. If *p*-value is small, then it can be concluded that the AUC-ROC is significantly different from 0.5 and that therefore there is evidence that the model actually discriminates between significant and non-significant results [89]. Typically, a threshold value (called significance level) of *p*-value < 0.05 is used. However, biomedicine studies often use more restrictive values like 0.005 [28] or even 0.001 [4].

### 4.4. Network exposition, visualization and interaction

Last but not least, at the end of the pipeline the results obtained should come out in a format that can be consumed by the audience (e.g. the scientific community, the media or even ourselves to inform the next iteration). One of the major advantages of disease networks is the intuitive access to the underlying complex interactions between diseases and other diseases, genes or drugs. Thus, publishing not only the data but also means to explore and exploit the network is key to ensure reproducibility and extensibility of the study [90]. Early studies lacked this option, although access to their data allowed the construction of visualization tools a posteriori. For example, Ramiro Gómez created an interactive view of the Human Disease Network proposed by Goh in 2007 using the graph visualization software Gephi^2^ and the original dataset from the study^3^. The same software was used by in 2014 and by Hoehndorf in 2015 to visualize the generated disease networks [39]. In both cases, a force-directed layout was used for the graph drawing [91].

Advances in network visualization tools have prompted the publication of network exploration systems associated with studies, being Cystoscape^4^ a remarkable example. Cytoscape provides basic functionality to layout and query the network; to visually integrate the network with expression profiles, phenotypes, and other molecular states; and to link the network to databases of functional annotations [92]. A number of studies have used Cytoscape as a basis to build and visualize their networks. For instance, Le et al. created HGPEC as an app for Cytoscape to predict novel disease-gene and disease-disease associations [93]. DisGeNet provided another app that allows to visualize, query and analyse a network representation of DisGeNET database [94]. Many other apps can be found in the Cytoscape app store^5^.

On their side, Himmelstein et al. accompanied their study based on heterogeneous disease networks with a powerful visualization tool built with Neo4j^6^ [46] that provides browsing and querying on Hetionet (see **Figure 5**). Being a remarkable example of data accessibility, not only the data but also the code of this tool is publicly available. Different studies of the University of Rome, such as SIGNOR^7^ and DISNOR^8^, also provides a disease network visualization tool that includes intuitive representations of the interactions between biological entities at different complexity levels (see **Figure 6**). This visualization tool was developed ad-hoc for these projects [95–97].

**Figure 5.**
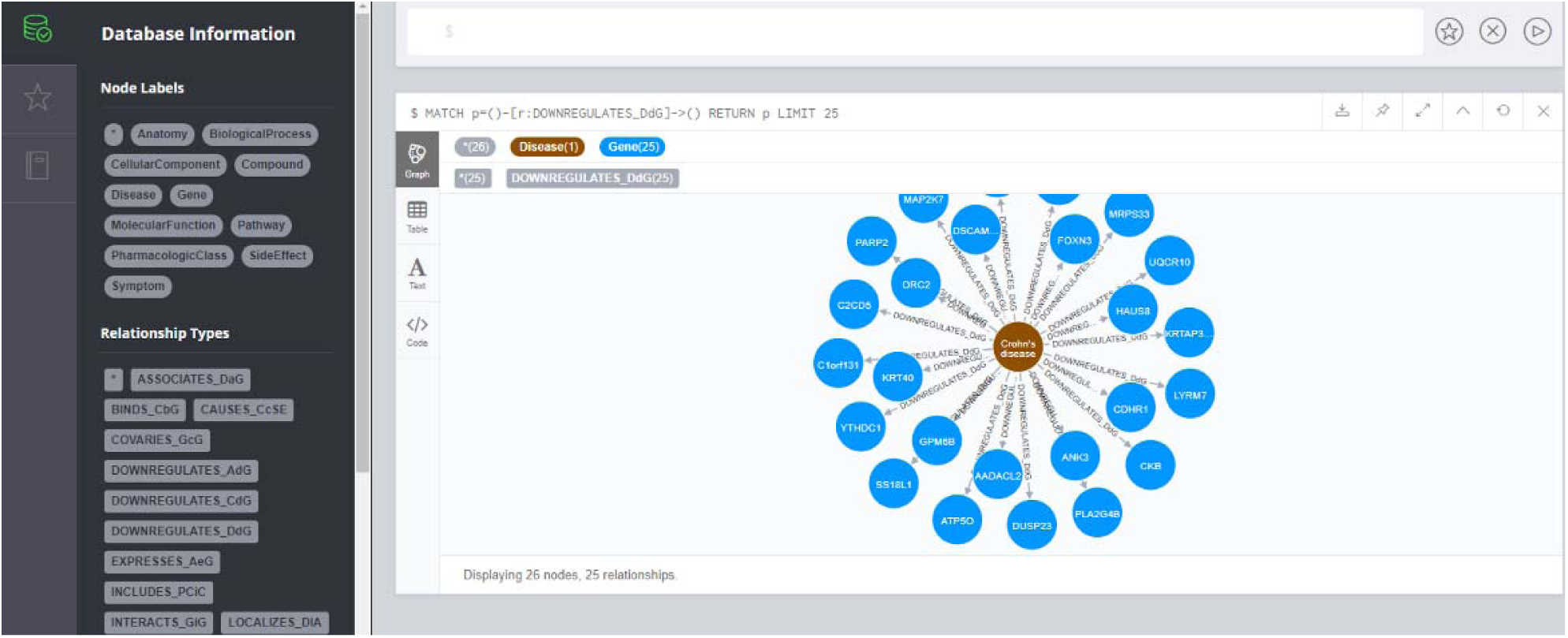
Hetionet Neo4j browser displaying semantic disease-gene associations for Crohn’s disease in an intearctive way. Retrieved from https://neo4j.het.io/browser. Copyright: 2015 Himmelstein, Baranzini. This is an open access project distributed under the terms of the Creative Commons Universal License.

**Figure 6.**
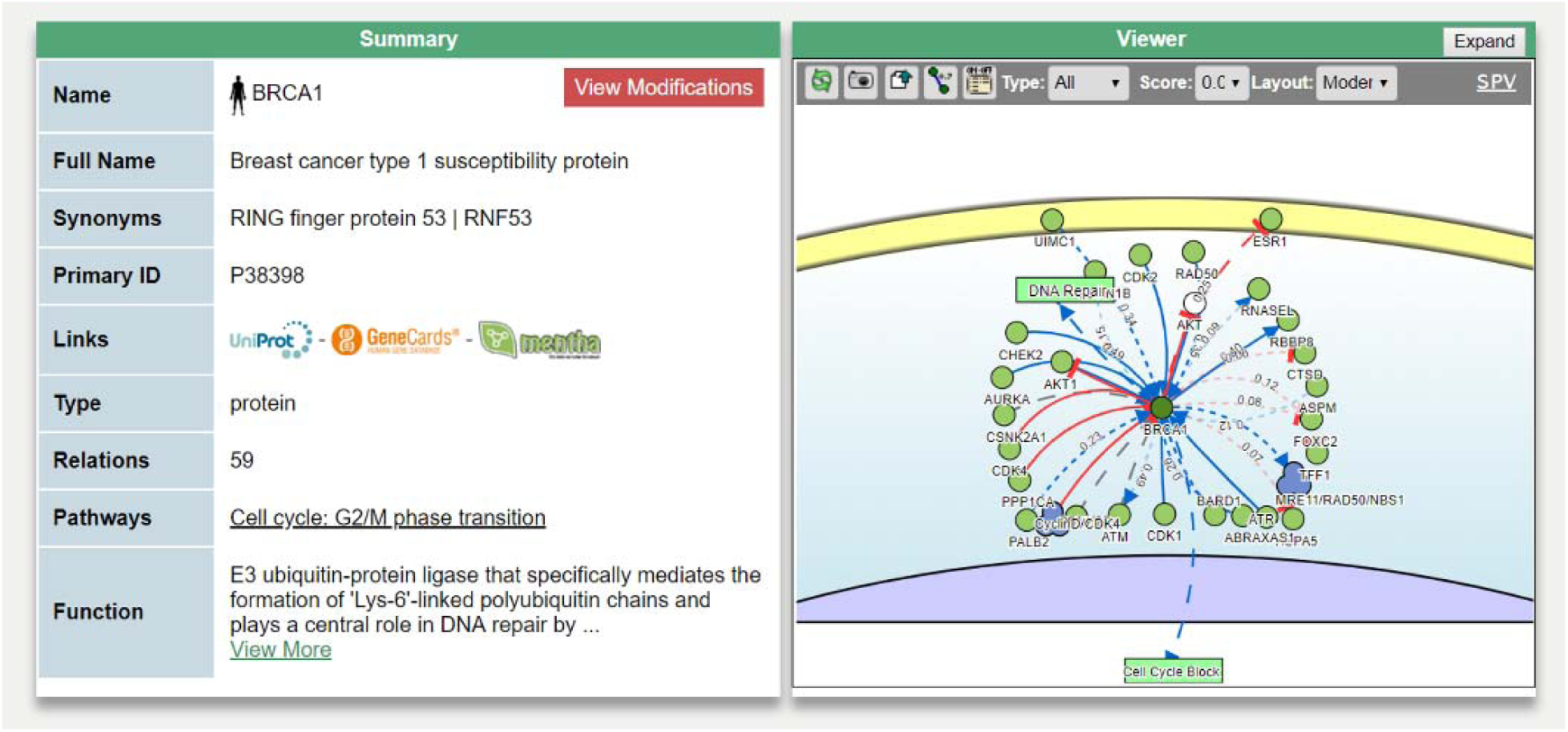
DISNOR signaling network browser showing an interactive graph for a protein related with Breast Cancer. Retrieved from https://disnor.uniroma2.it. Copyright: 2018, SIGNOR. This is an open access project distributed under the terms of the Creative Commons Attribution-ShareAlike 4.0 International License.

A recent study by Pavlopoulos et al. performs an empirical comparison of visualization tools for large-scale network analysis [98].

## 5. Discussion

The analysis of the evolution of the disease networks carried out in the first part of the document shows how these models have become increasingly complex and allow to address arduous problems such as the improvement of our disease understanding or the repositioning of drugs with promising results. However, as a side effect of this growing complexity, new challenges have emerged that need to be addressed.

The growing availability of biological sources, key in the improvement of disease networks, is ballasted by their fragmentation, heterogeneity, availability and different conceptualization of their data [3]. Furthermore, these sources are intrinsically biased towards consolidated knowledge, which complicates the discovery of novel findings. The exploitation of textual sources such as clinical histories or scientific articles – more abundant and faster growing – allows researchers to compensate for these limitations. As an example of the abundance and potential of these alternative sources, in a recent study Westergaard extracted and analyzed 15 million English scientific full-text articles published during the period 1823–2016 [99].

Despite this demonstrated potential, the exploitation of medical literature is not exempt from limitations. For instance, in the aforementioned study by Westergaard, the team could only access a subset of the Medline articles in full-text mode, while for the rest only the abstracts were available. In addition, depending on the source, they had to process documents with different structures and format. The use of open and structured sources such as Wikipedia has recently been proposed as an alternative [100]. Noise is also an important limiting factor when integrating new sources to enhance the predictive capacity of disease networks [101]. Adding new sources does not necessarily imply an improvement, since some databases are more informative than others. For example, Žitnik et al. evaluated the impact of removing sources in the performance of the proposed model to validate their informativeness. They observed that while the absence of some sources significantly affected the performance, in other case the impact was minimum [4]. It is therefore necessary to counteract this effect by choosing algorithms which can weigh the relevance and confidence of various sources of information (either within a single dataset, or across datasets) and discard those irrelevant before constructing the model [49].

Validation is yet another challenge in the studies based on disease networks. In some cases, the absence of a *gold standard* leads to the use of previous studies for the validation of the new models [28, 82], which might result in the propagation of errors from one study to another. The use of curated sources and of sufficiently contrasted studies helps to partially alleviate this problem [49, 73]. Nevertheless, in the case of drug repositioning, the validation of a drug’s ability to treat another disease ultimately requires in-vitro and in-vivo validations.

Related with the challenge of validation, the difficulty in accessing data from some studies prevents their reproducibility and verification by other teams, which makes them less reliable as references for future studies or as benchmarks. However, the effort of some researchers in making available the results of their work is worth to mention. Study cases such as Hetionet, Rephetio, SIGNOR and DisNOR [46, 47, 95, 96], which offer advanced search and visualization tools, undoubtedly represent the path to follow.

The review of the process of creating a disease network from the point of view of a data science pipeline carried out in the second part of the document allows to compare how each study has faced these challenges. **Supplementary Table 5** lists some of the most notable studies related to disease networks of the last decade, breaking down each of its phases. It also contains information on the type of problem addressed and the characteristics of the obtained network. This table could be considered an extension/update of the one compiled by Sun K. et al. [41].

## 6. Open questions and future lines of research

As a result of the systematized review of the literature on disease network reconstruction carried out in sections 2 and 3, and based on the previous discussion, we present in this section some of the open questions in this area of research and propose future lines of work to address them.

### 6.1. Source integration

Given the upwards trend to integrate new sources for the construction of more complete disease networks, a question that arises is *what sources remain to be added?*. In the case of biological sources, for example, while **Supplementary Table 1** contains some of the most commonly used genomic, metabolomic, proteomic, transcriptomic or phenomic resources, there are dozens of other ‘omics’ to explore [102], such as cytomics (cellular systems) or microbiomics (microbiota).

Further open questions related to the integration of sources have to do with the mechanisms used for their exploitation. Again, **Supplementary Tables 1-3** contain information about the form of access provided by each source, but *are there other services or alternative tools?* Their compilation and comparative analysis would allow completing the quick reference guide provided by these tables.

Source integration often involves mapping identifiers of diseases, genes or drugs, for example. **Supplementary Table 4** includes some of the sources commonly used for this purpose. However, the process is still tedious and prone to errors. *Are there tools that facilitate or even automate this task?* Research on the available solutions and the eventual development of a tool for this purpose are considered as possible lines of work.

Finally, as previously explained, the aggregation of sources to improve the completeness of disease networks has the introduction of noise as a collateral effect. *How can this effect be avoided?* In the study, some works addressing this problem have been presented, but given its importance, a more detailed research is required.

### 6.2. Methodology

Throughout our study, we reviewed the methods to build and analyze disease networks proposed by researchers over time. **Supplementary Table 5** outlines some of these studies. However, our analysis does not answer questions such as *what measure of similarity is more convenient in each case?* or *what network analysis methods allow better results in detecting disease similarities or repositioning drugs?* Therefore, a comparative study of the methodology of construction and analysis of disease networks would be a very valuable contribution to this field. Such study requires the use of a usefulness metric, or at least of a way of comparing similar results; yet, this is still not available, as discussed below.

### 6.3. Validation

The difficulty in validating the results of studies based on disease networks has been highlighted in previous sections. Given the variety of objectives (e.g. novel understanding, new drug applications), *what criteria should be used in each case to determine if a network is correct?* Moreover, while expert validation – as well as in vitro and in vivo tests – are still essential, the access to reliable sources or *gold standards* facilitates “in silico” analysis prior to these costly tests. *What are the “gold standards” available for each type of studie?* A compilation of these resources classified by study objective would be enormously useful in conducting new studies.

### 6.4. Visualization

One of the main advantages of the use of biological networks in the improvement of our disease understanding is their intuitiveness. While the graphic visualization of the disease network is just another form of representation (not to be confused with the network itself), it facilitates its interpretation. However, as described in section 4.4, the lack of a standard leads to different studies and different tools generating different visualizations, which may bias this interpretation.. The lines of work in this area should be oriented to answer the questions *What rules can be proposed to alleviate these problems when representing disease networks graphically? Could a standard be proposed?*

## 7. Conclusion

Research studies on based disease networks have significantly advanced over the last decade. From the initial simple undirected networks that associated diseases with symptoms or genes in a way, we have moved to complex networks that relate the disease to dozens of features from different sources in a semantic, directional and weighted way. The growing availability of biological and textual sources, the improvement in techniques and processing capacity and the use of new models have contributed fundamentally to this progress. As can be concluded from the analysis in the first part of the document, the contribution of disease networks to fields of disease understanding and drug repositioning is increasingly notable.

Nevertheless, an exhaustive analysis of the phases in the process of creating disease networks carried out in the second half of the document reveals important challenges. First, biological sources suffer from fragmentation, heterogeneity, lack of availability and different conceptualization, that can only be alleviated in part with the aggregation of textual sources. Second, the combination of sources involves the introduction of noise that can affect the performance of the model, which makes it necessary to take preventive measures in this regard. Finally, the scarcity of reference data and verifiable studies hinders the validation of the new models.

In addition to detecting these challenges, the analysis of disease networks from the point of view of their functional units allows for a more precise comparison of studies, highlighting their differences and common points. This study and the presented analyses, reflected in the summary tables, can serve to inspire future work. On one hand, a performance comparison of the prediction models accross the different studies might lead to deduce which functional units offer better results. On the other hand, the analysis of these units can facilitate the detection of unexploited sources or methods and explore them as alternatives. In a next phase, based on the obtained results, alternative combinations of these functional units could be proposed to build new pipelines and obtain more precise models based on disease networks.

---

**Supplementary Table 1.**
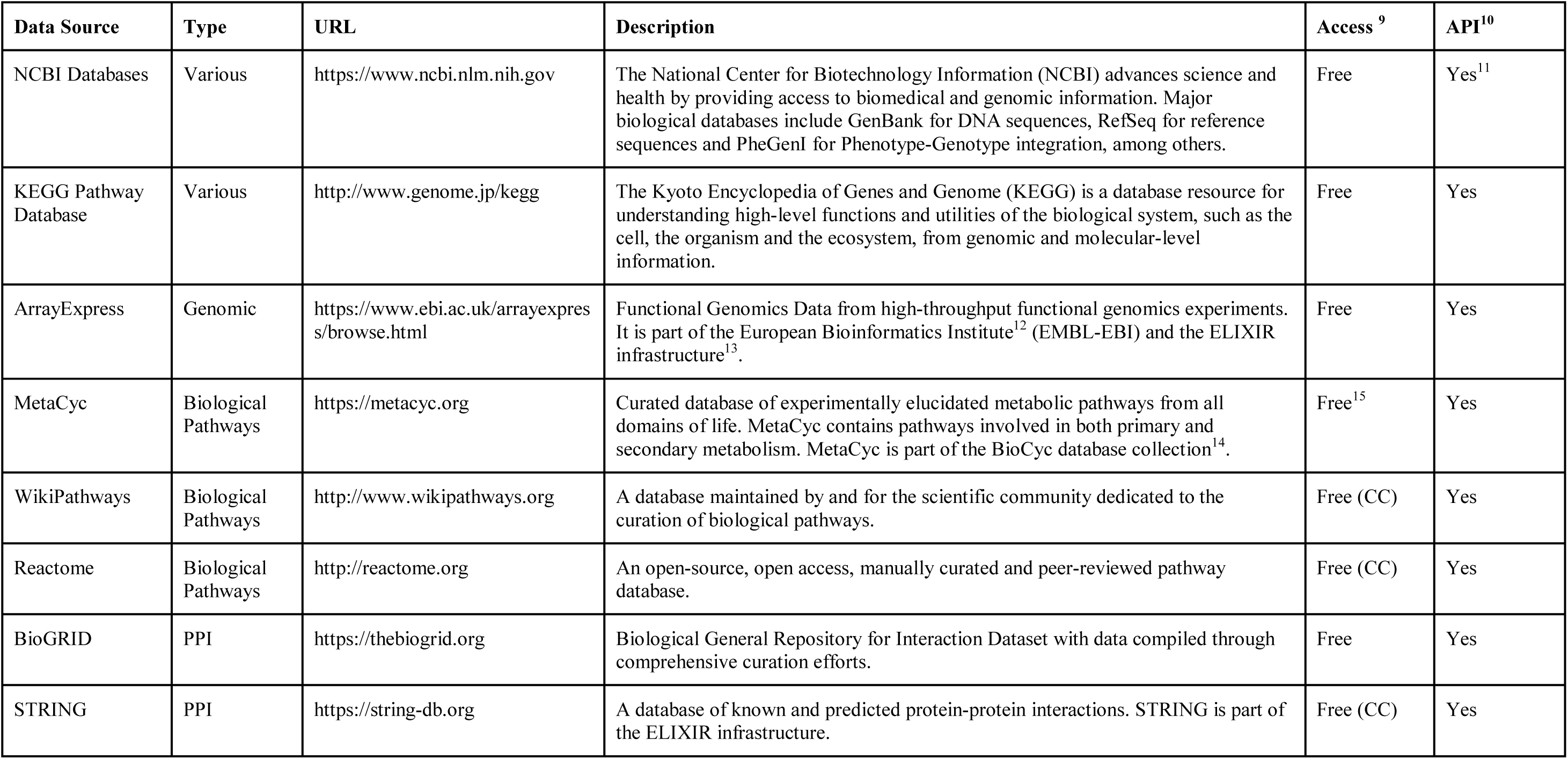
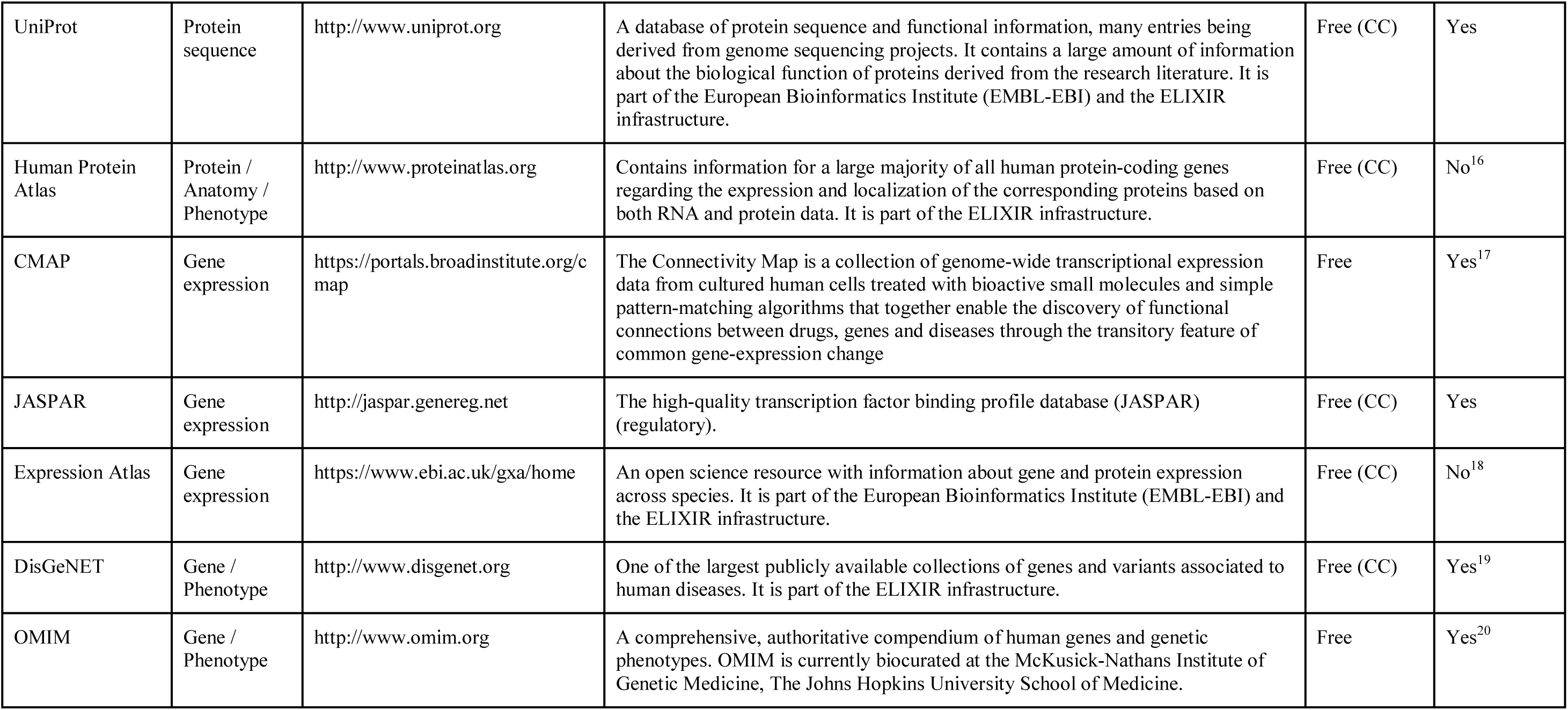
Biological data sources.

**Supplementary Table 2.** Drug data sources.

| Data Source | Type | URL | Description | Access | API |
| --- | --- | --- | --- | --- | --- |
| FDA Databases | Various | https://www.fda.gov/Drugs/InformationOnDrugs | Information about FDA-approved brand name and generic prescription and over-the-counter human drugs and biological therapeutic products. | Free | Yes<sup>21</sup> |
| PubChem | Compound | https://pubchem.ncbi.nlm.nih.gov | A database of chemical molecules and their activities against biological assays. The system is maintained by the National Center for Biotechnology Information (NCBI), a component of the National Library of Medicine, which is part of the United States National Institutes of Health (NIH). | Free | Yes<sup>22</sup> |
| DrugBank | Phenotype | http://www.drugbank.ca | This database combines detailed drug (i.e. chemical, pharmacological and pharmaceutical) data with comprehensive drug target (i.e. sequence, structure, and pathway) information. | Free (CC) | Yes<sup>23</sup> |
| Orphanet | Phenotype | http://www.orphadata.org | A unique resource, gathering and improving knowledge on rare diseases so as to improve the diagnosis, care and treatment of patients with rare diseases. The Orphanet Rare Disease ontology (ORDO) provides a structured vocabulary for rare diseases capturing relationships between diseases, genes and other relevant features. | Free (CC) | No<sup>24</sup> |
| SIDER | Phenotype | http://sideeffects.embl.de | Contains information on marketed medicines and their recorded adverse drug reactions. The information is extracted from public documents and package inserts. The available information includes side effect frequency, drug and side effect classifications as well as links to further information, for example drug–target relations. | Free (CC) | No<sup>25</sup> |
| PharmaGKB | Gene / Phenotype | http://www.pharmgkb.org | Pharmacogenomics knowledge resource that encompasses clinical information including dosing guidelines and drug labels, potentially clinically actionable gene-drug associations and genotype-phenotype relationships | Free (CC) | Yes<sup>26</sup> |
| ChEMBL | Compound | https://www.ebi.ac.uk/chembl | Database of bioactive drug-like small molecules, it contains 2-D structures, calculated properties and abstracted bioactivities. It is part of the European Bioinformatics Institute (EMBL-EBI) and the ELIXIR infrastructure. | Free | Yes |

<sup>21</sup> Open FDA Access Key is needed to get the maximum rate of requests per minute. https://open.fda.gov/apis/authentication/
<sup>22</sup> Limited to 5 requests per second. https://pubchemdocs.ncbi.nlm.nih.gov/pug-rest
<sup>23</sup> A paid subscription is required to access the API. Data is available through online search and downloads.
<sup>24</sup> Downloads available. SPARQL endpoint available to consume the ontology.
<sup>25</sup> Data are available via online search and downloads

|  |  |  |  |  |  |
| --- | --- | --- | --- | --- | --- |
| Drug Repurposing Hub | Compound | https://clue.io/repurposing | Contains extensive curated annotations for each drug, including details about commercial sources of all compounds | Free | Yes<sup>27</sup> |

**Supplementary Table 3.** Medical textual sources.

| Data Source | Type | URL | Description | Access | API |
| --- | --- | --- | --- | --- | --- |
| PubMed/PMC | Journals | https://www.nlm.nih.gov/bsd/pmresources.html | PubMed contains more than 28 million medical journal citations from MEDLINE indexed journals and NCI Bookshelf. These citations may have links to full-text articles or manuscripts in PMC (PubMed Central). | Free | Yes<sup>28</sup> |
| MedlinePlus | Medical Encyclopedia | https://medlineplus.gov | Provides encyclopedic information on health and drug issues, and provides a directory of medical services. The service provides curated consumer health information in English and Spanish. | Free | Yes |
| ClinicalTrials | Clinical Trials | http://www.clinicaltrials.gov | A database of privately and publicly funded clinical studies conducted around the world. It is a resource provided by the U.S. National Library of Medicine. | Free | No<sup>29</sup> |
| Europe PMC | Journals / Clinical Trials | http://europepmc.org | Provides access to worldwide life sciences articles, books, patents and clinical guidelines. Europe PMC provides links to relevant records in databases such as Uniprot, European Nucleotide Archive (ENA), Protein Data Bank Europe (PDBE) and BioStodie. It is part of the ELIXIR infrastructure. | Free | Yes |
| GWAS Catalog | GWAS Studies | https://www.ebi.ac.uk/gwas/ | The NHGRI-EBI Catalog of published genome-wide association studies. Eligible studies are curated within 1-2 months of publication, dependent on the availability of literature, and the data is released on a weekly cycle | Free | Yes |

<sup>28</sup> Entrez: API key required (as of May 1, 2018) to reach a request rate up to 10 per second. See https://www.ncbi.nlm.nih.gov/books/NBK25500.
<sup>29</sup> Not strictly an API. The combination of Advanced Search query building and results download provides intensive data access.

**Table 4.** Mapping sources.

| Data Source | Type | URL | Description | Access | API |
| --- | --- | --- | --- | --- | --- |
| MeSH | Medical Thesaurus | https://www.ncbi.nlm.nih.gov/mesh | Medical Subject Headings (MeSH) is the NLM controlled vocabulary thesaurus used for indexing articles for PubMed. It is also used by ClinicalTrials.gov registry to classify which diseases are studied by trials. | Free | Yes<sup>30</sup> |
| SNOMED CT | Medical Thesaurus | http://www.snomed.org/snomed-ct | Systematized Nomenclature of Medicine Clinical Terms (SNOMED CT) is a comprehensive and precise clinical health terminology product. | Licensed<sup>31</sup> | Yes |
| UMLS | Medical Thesaurus | https://www.nlm.nih.gov/research/umls/ | Unified Medical Language System (UMLS) integrates and distributes key terminology, classification and coding standards, and associated resources to promote creation of more effective and interoperable biomedical information systems and services, including electronic health records | Licensed<sup>32</sup> | Yes |
| ICD-CM | Disease Codes | http://www.who.int/classifications/icd | International Classification of Diseases (ICD) is the classification used to code and classify mortality data from death certificates. ICD-Clinical Modification (CM) is used to code and classify morbidity data from the inpatient and outpatient records, physician offices, and most National Center for Health Statistics (NCHS) surveys. | Free | Yes<sup>33</sup> |
| HGNC | Gene Codes | https://www.genenames.org/ | HGNC is responsible for approving unique symbols and names for human loci, including protein coding genes, ncRNA genes and pseudogenes, to allow unambiguous scientific communication. | Free | Yes |
| DO | Disease Ontology | https://bioportal.biontology.org/ontologies/DOID | The Disease Ontology (DO) provides the biomedical community with consistent, reusable and sustainable descriptions of human disease terms, phenotype characteristics and related medical vocabulary disease concepts. The DO semantically integrates disease and medical vocabularies through extensive cross mapping of DO terms to MeSH, ICD, NCI's thesaurus, SNOMED and OMIM. | Free (CC) | Yes |
| PO | Phenotype Ontology | https://hpo.jax.org/app/ | The Human Phenotype Ontology (HPO) provides a standardized vocabulary of phenotypic abnormalities encountered in human disease. The HPO is currently being developed using the medical literature, Orphanet, DECIPHER, and OMIM. | Free | Yes<sup>34</sup> |
| GO | Gene Ontology | http://www.geneontology.org | The Gene Ontology (GO) provides a computational representation of our evolving | Free (CC) | Yes |

<sup>30</sup> API-like access through the SPARQL endpoint. https://hhs.github.io/meshrdf/sparql-and-uri-requests
<sup>31</sup> Free license for member countries. See other fee exemptions: https://www.snomed.org/snomed-ct/get-snomed-ct
<sup>32</sup> Free license available under certain conditions. https://uts.nlm.nih.gov/license.html
<sup>33</sup> Provided by NLM Clinical Tables Search Service: https://clinicaltables.nlm.nih.gov/apidoc/icd10cm/v3/doc.html
<sup>34</sup> SPARQL endpoint available at http://www.orphadata.org/cgi-bin/inc/sparql_hoom.inc.php

|  |  |  |  |  |  |
| --- | --- | --- | --- | --- | --- |
|  |  | logy.org | knowledge of how genes encode biological functions at the molecular, cellular and tissue system levels. GO terms are organised in three domains: cellular component, molecular function and biological process. |  |  |
| Uberon | Anatomy Ontology | http://uberon.github.io/ | An integrated cross-species ontology covering anatomical structures in animals. See the about page for more info, or read the Uberon paper in Genome Biology | Free | Yes<sup>35</sup> |

**Supplementary Table 5.**
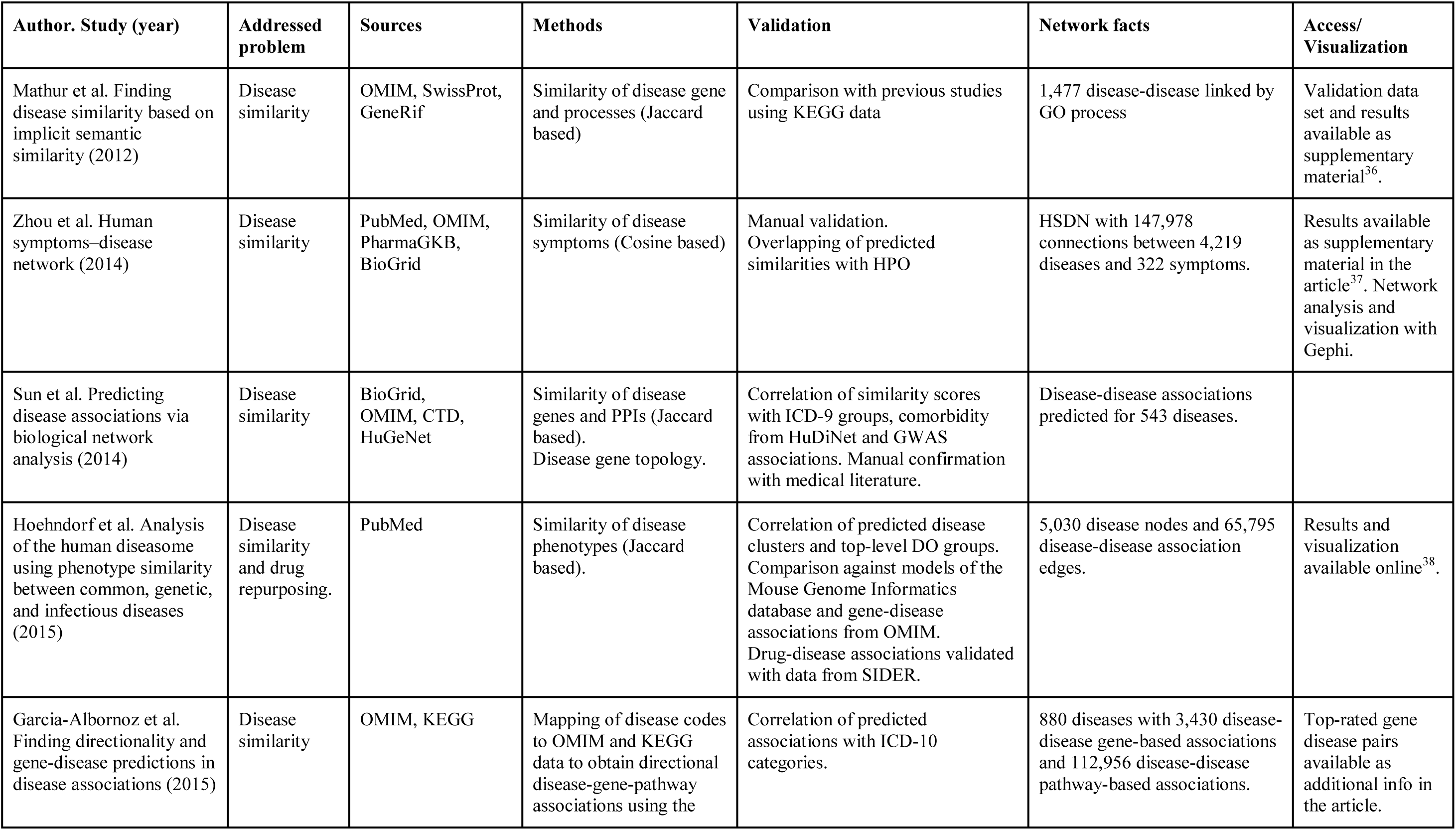
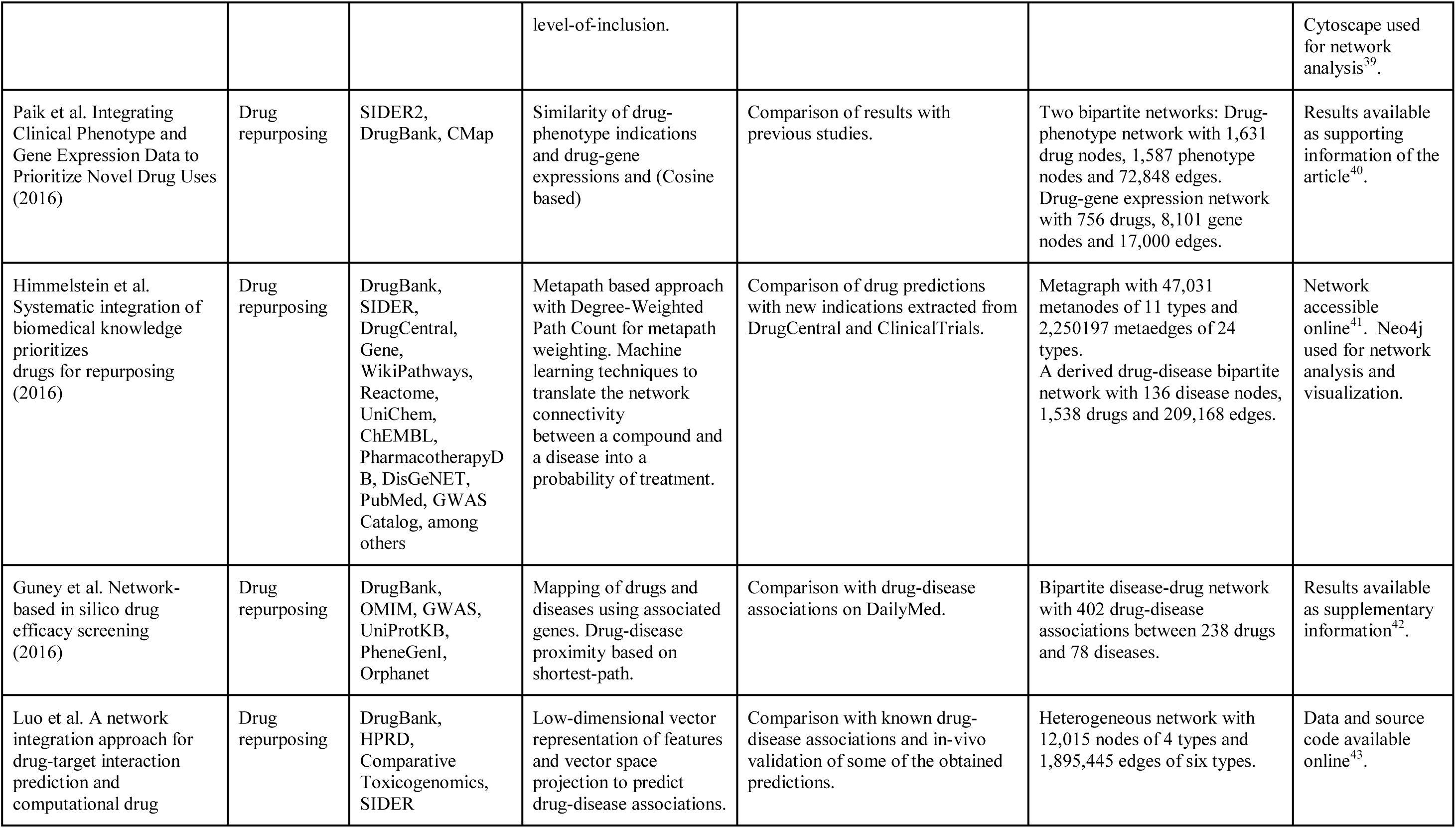
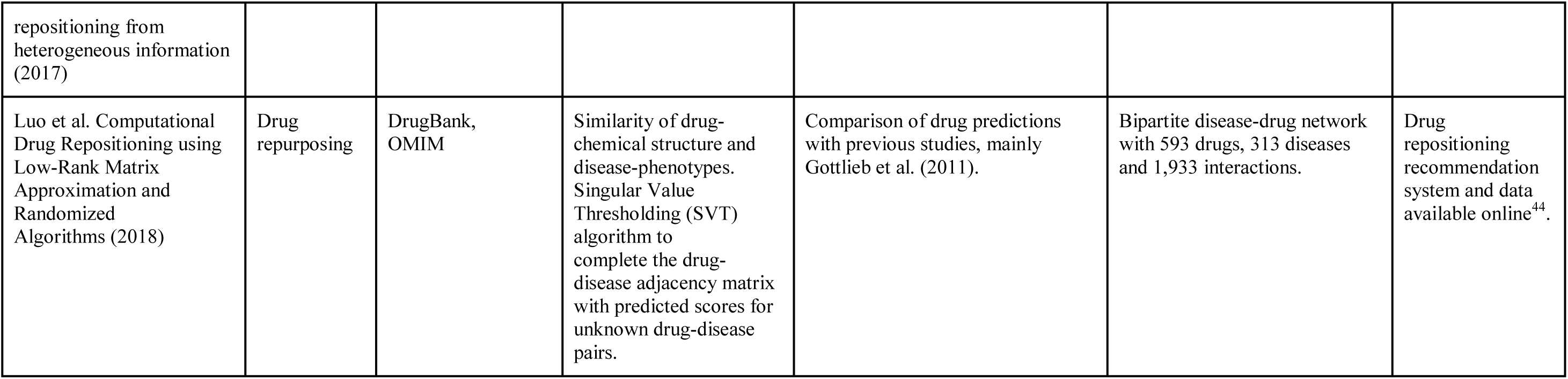
Studies on disease networks.

## Footnotes

1 https://biopython.org

2 https://gephi.org

3 https://exploring-data.com/info/human-disease-network

4 http://www.cytoscape.org

5 http://apps.cytoscape.org

6 https://neo4j.com

7 https://signor.uniroma2.it

8 https://disnor.uniroma2.it

9 To resouces, online search, downloads and/or API, least for academic purposes. Databases with a Creative Commons License type are marked with CC. For commercial use, please refer to licensing details on the Databse URL.

10 The database provides a consumible API for extensive use, via tools and/or web services. See details on database URL.

11 Entrez: API key required (as of May 1, 2018) to reach a request rate up to 10 per second. See https://www.ncbi.nlm.nih.gov/books/NBK25500.

12 https://www.ebi.ac.uk/

13 https://www.elixir-europe.org/

14 https://biocyc.org

15 Access via BioCyc web services require paid subscription, but MetaCyc search online and downloads are free access.

16 Data is accessible via downloads. A programmatic access to filter downloadable data is provided. https://www.proteinatlas.org/about/help/dataaccess

17 Clue API Key required for unlimited use. https://clue.io/api

18 Tools for R available, but data must be downloaded first.

19 API-like access through SPARQL endpoint http://rdf.disgenet.org/sparql/

20 Registration is required. https://omim.org/api

21 Open FDA Access Key is needed to get the maximum rate of requests per minute. https://open.fda.gov/apis/authentication/

22 Limited to 5 requests per second. https://pubchemdocs.ncbi.nlm.nih.gov/pug-rest

23 A paid subscription is required to access the API. Data is available through online search and downloads.

24 Downloads available. SPARQL endpoint available to consume the ontology.

25 Data are available via online search and downloads

26 Beta version. Limited to 2 requests per second. https://api.pharmgkb.org/

27 Clue API Key required for unlimited use. https://clue.io/api

28 Entrez: API key required (as of May 1, 2018) to reach a request rate up to 10 per second. See https://www.ncbi.nlm.nih.gov/books/NBK25500.

29 Not strictly an API. The combination of Advanced Search query building and results download provides intensive data access.

30 API-like access through the SPARQL endpoint. https://hhs.github.io/meshrdf/sparql-and-uri-requests

31 Free license for member countries. See other fee exemptions: https://www.snomed.org/snomed-ct/get-snomed-ct

32 Free license available under certain conditions. https://uts.nlm.nih.gov//license.html

33 Provided by NLM Clinical Tables Search Service: https://clinicaltables.nlm.nih.gov/apidoc/icd10cm/v3/doc.html

34 SPARQL endpoint available at http://www.orphadata.org/cgi-bin/inc/sparql_hoom.inc.php

35 SPARQL endopoint available: http://sparql.hegroup.org/sparql

36 https://ars.els-cdn.com/content/image/1-s2.0-S1532046411002073-mmc1.doc

37 https://www.nature.com/articles/ncomms5212#s1

38 http://aber-owl.net/aber-owl/diseasephenotypes

39 https://static-content.springer.com/esm/art%3A10.1186%2Fs12918-015-0184-9/MediaObjects/12918_2015_184_MOESM1_ESM.xlsx

40 https://ascpt.onlinelibrary.wiley.com/doi/full/10.1002/psp4.12108

41 http://het.io

42 https://www.ncbi.nlm.nih.gov/pmc/articles/PMC4740350/#S1

43 https://github.com/luoyunan/DTINet

44 http://bioinformatics.csu.edu.cn/resources/softs/DrugRepositioning/DRRS/index.html

## References

1. Goh K-I, Cusick ME, Valle D, Childs B, Vidal M, Barabási A-L. The human disease network. PNAS. 2007;104:8685–90.

2. Yang J, Wu S-J, Dai W-T, Li Y-X, Li Y-Y. The human disease network in terms of dysfunctional regulatory mechanisms. Biol Direct. 2015;10. doi:10.1186/s13062-015-0088-z.

3. Loscalzo J, Kohane I, Barabasi A-L. Human disease classification in the postgenomic era: A complex systems approach to human pathobiology. Mol Syst Biol. 2007;3:124.

4. Žitnik M, Janjić V, Larminie C, Zupan B, Pržulj N. Discovering disease-disease associations by fusing systems-level molecular data. Scientific Reports. 2013;3:3202.

5. Chan SY, Loscalzo J. The emerging paradigm of network medicine in the study of human disease. Circ Res. 2012;111:359–74.

6. Strogatz SH. Exploring complex networks. Nature. 2001;410:268–76.

7. Albert R, Barabási A-L. Statistical mechanics of complex networks. Rev Mod Phys. 2002;74:47–97.

8. Newman M. The Structure and Function of Complex Networks. SIAM Rev. 2003;45:167–256.

9. Boccaletti S, Latora V, Moreno Y, Chavez M, Hwang D-U. Complex networks: Structure and dynamics. Physics Reports. 2006;424:175–308.

10. Costa L da F, Rodrigues FA, Travieso G, Boas PRV. Characterization of complex networks: A survey of measurements. Advances in Physics. 2007;56:167–242.

11. Barabási A-L. Network Medicine — From Obesity to the “Diseasome.” New England Journal of Medicine. 2007;357:404–7.

12. Vocaturo E, Veltri P. On the use of Networks in Biomedicine. Procedia Computer Science. 2017;110:498–503.

13. Park S, Lee D, Shin H. Network mirroring for drug repositioning. BMC Med Inform Decis Mak. 2017;17 Suppl 1. doi:10.1186/s12911-017-0449-x.

14. Grant MJ, Booth A. A typology of reviews: an analysis of 14 review types and associated methodologies. Health Info Libr J. 2009;26:91–108.

15. Goh K-I, Choi I-G. Exploring the human diseasome: the human disease network. Brief Funct Genomics. 2012;11:533–42.

16. Lee D-S, Park J, Kay KA, Christakis NA, Oltvai ZN, Barabási A-L. The implications of human metabolic network topology for disease comorbidity. PNAS. 2008;105:9880–5.

17. Barrenas F, Chavali S, Holme P, Mobini R, Benson M. Network Properties of Complex Human Disease Genes Identified through Genome-Wide Association Studies. PLoS One. 2009;4. doi:10.1371/journal.pone.0008090.

18. Mullen J, Cockell SJ, Woollard P, Wipat A. An Integrated Data Driven Approach to Drug Repositioning Using Gene-Disease Associations. PLoS One. 2016;11. doi:10.1371/journal.pone.0155811.

19. Hernandez JJ, Pryszlak M, Smith L, Yanchus C, Kurji N, Shahani VM, et al. Giving Drugs a Second Chance: Overcoming Regulatory and Financial Hurdles in Repurposing Approved Drugs As Cancer Therapeutics. Front Oncol. 2017;7:273.

20. Altshuler D, Daly M, Kruglyak L. Guilt by association. Nat Genet. 2000;26:135–7.

21. Yildirim MA, Goh K-I, Cusick ME, Barabási A-L, Vidal M. Drug-target network. Nat Biotechnol. 2007;25:1119–26.

22. Ma’ayan A, Jenkins SL, Goldfarb J, Iyengar R. Network Analysis of FDA Approved Drugs and their Targets. Mt Sinai J Med. 2007;74:27–32.

23. Chiang AP, Butte AJ. Systematic evaluation of drug-disease relationships to identify leads for novel drug uses. Clin Pharmacol Ther. 2009;86:507–10.

24. Bleakley K, Yamanishi Y. Supervised prediction of drug–target interactions using bipartite local models. Bioinformatics. 2009;25:2397–403.

25. Campillos M, Kuhn M, Gavin A-C, Jensen LJ, Bork P. Drug target identification using side-effect similarity. Science. 2008;321:263–6.

26. Hu G, Agarwal P. Human Disease-Drug Network Based on Genomic Expression Profiles. PLoS One. 2009;4. doi:10.1371/journal.pone.0006536.

27. Suthram S, Dudley JT, Chiang AP, Chen R, Hastie TJ, Butte AJ. Network-based elucidation of human disease similarities reveals common functional modules enriched for pluripotent drug targets. PLoS Comput Biol. 2010;6:e1000662.

28. Mathur S, Dinakarpandian D. Finding disease similarity based on implicit semantic similarity. J Biomed Inform. 2012;45:363–71.

29. Rzhetsky A, Wajngurt D, Park N, Zheng T. Probing genetic overlap among complex human phenotypes. PNAS. 2007;104:11694–9.

30. Hidalgo CA, Blumm N, Barabási A-L, Christakis NA. A Dynamic Network Approach for the Study of Human Phenotypes. PLOS Computational Biology. 2009;5:e1000353.

31. Jiang Y, Ma S, Shia B-C, Lee T-S. An Epidemiological Human Disease Network Derived from Disease Co-occurrence in Taiwan. Scientific Reports. 2018;8:4557.

32. Conway M, Berg RL, Carrell D, Denny JC, Kho AN, Kullo IJ, et al. Analyzing the heterogeneity and complexity of Electronic Health Record oriented phenotyping algorithms. AMIA Annu Symp Proc. 2011;2011:274–83.

33. Yip C, Han N-LR, Sng BL. Legal and ethical issues in research. Indian J Anaesth. 2016;60:684–8.

34. Okumura T, Aramaki E, Tateisi Y. Clinical Vocabulary and Clinical Finding Concepts in Medical Literature. In: The First Workshop on Natural Language Processing for Medical and Healthcare Fields. Nagoya: Asian Federation of Natural Language Processing; 2013. p. 7–13. http://www.aclweb.org/anthology/W13-4602. Accessed 3 Sep 2018.

35. Rodríguez-González A, Martínez-Romero M, Costumero R, Wilkinson MD, Menasalvas-Ruiz E. Diagnostic Knowledge Extraction from MedlinePlus: An Application for Infectious Diseases. In: Overbeek R, Rocha MP, Fdez-Riverola F, De Paz JF, editors. 9th International Conference on Practical Applications of Computational Biology and Bioinformatics. Springer International Publishing; 2015. p. 79–87.

36. Rodríguez González A, Costumero Moreno R, Martínez Romero M, Wilkinson MD, Menasalvas Ruiz E. Extracting diagnostic knowledge from MedLine Plus: a comparison between MetaMap and cTAKES Approaches. Current Bioinformatics. 2015;375:1–7.

37. Rebholz-Schuhmann D, Oellrich A, Hoehndorf R. Text-mining solutions for biomedical research: enabling integrative biology. Nat Rev Genet. 2012;13:829–39.

38. Zhou X, Menche J, Barabási A-L, Sharma A. Human symptoms–disease network. Nature Communications. 2014;5:4212.

39. Hoehndorf R, Schofield PN, Gkoutos GV. Analysis of the human diseasome using phenotype similarity between common, genetic, and infectious diseases. Sci Rep. 2015;5:10888.

40. Gottlieb A, Stein GY, Ruppin E, Sharan R. PREDICT: a method for inferring novel drug indications with application to personalized medicine. Mol Syst Biol. 2011;7:496.

41. Sun K, Gonçalves JP, Larminie C, Pržulj N. Predicting disease associations via biological network analysis. BMC Bioinformatics. 2014;15:304.

42. Garcia-Albornoz M, Nielsen J. Finding directionality and gene-disease predictions in disease associations. BMC Syst Biol. 2015;9. doi:10.1186/s12918-015-0184-9.

43. Daminelli S, Haupt VJ, Reimann M, Schroeder M. Drug repositioning through incomplete bi-cliques in an integrated drug-target-disease network. Integr Biol (Camb). 2012;4:778–88.

44. Wang W, Yang S, Zhang X, Li J. Drug repositioning by integrating target information through a heterogeneous network model. Bioinformatics. 2014;30:2923–30.

45. Chen B, Ding Y, Wild DJ. Assessing Drug Target Association Using Semantic Linked Data. PLoS Comput Biol. 2012;8. doi:10.1371/journal.pcbi.1002574.

46. Himmelstein DS, Lizee A, Hessler C, Brueggeman L, Chen SL, Hadley D, et al. Systematic integration of biomedical knowledge prioritizes drugs for repurposing. eLife. 6. doi:10.7554/eLife.26726.

47. Himmelstein D, Lizee A, Hessler C, Brueggeman L, Chen S, Hadley D, et al. Rephetio: Repurposing drugs on a hetnet [report]. Thinklab. 2016. doi:10.15363/thinklab.a7.

48. Sun Y, Barber R, Gupta M, Aggarwal CC, Han J. Co-author Relationship Prediction in Heterogeneous Bibliographic Networks. In: 2011 International Conference on Advances in Social Networks Analysis and Mining. 2011. p. 121–8.

49. Luo Y, Zhao X, Zhou J, Yang J, Zhang Y, Kuang W, et al. A Network Integration Approach for Drug-Target Interaction Prediction and Computational Drug Repositioning from Heterogeneous Information. bioRxiv. 2017;:100305.

50. Ojeda T, Murphy SP, Bengfort B, Dasgupta A. Practical Data Science Cookbook. Packt Publishing; 2014.

51. Lu M, Zhang Q, Deng M, Miao J, Guo Y, Gao W, et al. An Analysis of Human MicroRNA and Disease Associations. PLOS ONE. 2008;3:e3420.

52. Zhou X, Lei L, Liu J, Halu A, Zhang Y, Li B, et al. A Systems Approach to Refine Disease Taxonomy by Integrating Phenotypic and Molecular Networks. EBioMedicine. 2018;31:79–91.

53. Gligorijević V, Pržulj N. Methods for biological data integration: perspectives and challenges. Journal of The Royal Society Interface. 2015;12:20150571.

54. Fan Y, Li M, Zhang P, Wu J, Di Z. The effect of weight on community structure of networks. Physica A: Statistical Mechanics and its Applications. 2007;378:583–90.

55. Zhou T, Ren J, Medo M, Zhang Y-C. Bipartite network projection and personal recommendation. Phys Rev E Stat Nonlin Soft Matter Phys. 2007;76 4 Pt 2:046115.

56. Salton G, Lesk ME. Computer Evaluation of Indexing and Text Processing. J ACM. 1968;15:8–36.

57. van Driel MA, Bruggeman J, Vriend G, Brunner HG, Leunissen JAM. A text-mining analysis of the human phenome. Eur J Hum Genet. 2006;14:535–42.

58. Sun K, Buchan N, Larminie C, Pržulj N. The integrated disease network. Integr Biol (Camb). 2014;6:1069–79.

59. Resnik P. Using Information Content to Evaluate Semantic Similarity in a Taxonomy. In: Proceedings of the 14th International Joint Conference on Artificial Intelligence – Volume 1. San Francisco, CA, USA: Morgan Kaufmann Publishers Inc.; 1995. p. 448–453. http://dl.acm.org/citation.cfm?id=1625855.1625914. Accessed 3 Sep 2018.

60. Lin D. An Information-Theoretic Definition of Similarity. In: In Proceedings of the 15th International Conference on Machine Learning. Morgan Kaufmann; 1998. p. 296–304.

61. Wang JZ, Du Z, Payattakool R, Yu PS, Chen C-F. A new method to measure the semantic similarity of GO terms. Bioinformatics. 2007;23:1274–81.

62. Cheng L, Li J, Ju P, Peng J, Wang Y. SemFunSim: A New Method for Measuring Disease Similarity by Integrating Semantic and Gene Functional Association. PLOS ONE. 2014;9:e99415.

63. Cheng L, Jiang Y, Wang Z, Shi H, Sun J, Yang H, et al. DisSim: an online system for exploring significant similar diseases and exhibiting potential therapeutic drugs. Scientific Reports. 2016;6:30024.

64. Hu Y, Zhao L, Liu Z, Ju H, Shi H, Xu P, et al. DisSetSim: an online system for calculating similarity between disease sets. J Biomed Semantics. 2017;8 Suppl 1. doi:10.1186/s13326-017-0140-2.

65. Omura M, Tateishi Y, Okumura T. Disease Similarity Calculation on Simplified Disease Knowledge Base for Clinical Decision Support Systems.: 6.

66. Dai W, Liu X, Gao Y, Chen L, Song J, Chen D, et al. Matrix Factorization-Based Prediction of Novel Drug Indications by Integrating Genomic Space. Computational and Mathematical Methods in Medicine. 2015. doi:10.1155/2015/275045.

67. Zhang W, Yue X, Lin W, Wu W, Liu R, Huang F, et al. Predicting drug-disease associations by using similarity constrained matrix factorization. BMC Bioinformatics. 2018;19:233.

68. Chen X, Liu M-X, Yan G-Y. Drug-target interaction prediction by random walk on the heterogeneous network. Mol Biosyst. 2012;8:1970–8.

69. Xie M, Hwang T, Kuang R. Prioritizing Disease Genes by Bi-Random Walk. In: Tan P-N, Chawla S, Ho CK, Bailey J, editors. Advances in Knowledge Discovery and Data Mining. Springer Berlin Heidelberg; 2012. p. 292–303.

70. Liu Y, Zeng X, He Z, Zou Q. Inferring MicroRNA-Disease Associations by Random Walk on a Heterogeneous Network with Multiple Data Sources. IEEE/ACM Trans Comput Biol Bioinformatics. 2017;14:905–915.

71. Yu G, Fu G, Lu C, Ren Y, Wang J. BRWLDA: bi-random walks for predicting lncRNA-disease associations. Oncotarget. 2017;8:60429–46.

72. Yang A, Troup M, Ho JWK. Scalability and Validation of Big Data Bioinformatics Software. Comput Struct Biotechnol J. 2017;15:379–86.

73. Jodeleit H, Palamides P, Beigel F, Mueller T, Wolf E, Siebeck M, et al. Design and validation of a disease network of inflammatory processes in the NSG-UC mouse model. Journal of Translational Medicine. 2017;15:265.

74. Seok J, Warren HS, Cuenca AG, Mindrinos MN, Baker HV, Xu W, et al. Genomic responses in mouse models poorly mimic human inflammatory diseases. PNAS. 2013;110:3507–12.

75. Perlman RL. Mouse models of human disease. Evol Med Public Health. 2016;2016:170–6.

76. Versi E. “Gold standard” is an appropriate term. BMJ. 1992;305:187–187.

77. D S, S S, A M. Measurement error correction for logistic regression models with an “alloyed gold standard”. American Journal of Epidemiology. 1997;145:184–96.

78. Kumar S, Chowdhury S, Kumar S. In silico repurposing of antipsychotic drugs for Alzheimer’s disease. BMC Neurosci. 2017;18. doi:10.1186/s12868-017-0394-8.

79. Suratanee A, Plaimas K. Network-based association analysis to infer new disease-gene relationships using large-scale protein interactions. PLOS ONE. 2018;13:e0199435.

80. Piñero J, Bravo À, Queralt-Rosinach N, Gutiérrez-Sacristán A, Deu-Pons J, Centeno E, et al. DisGeNET: a comprehensive platform integrating information on human disease-associated genes and variants. Nucleic Acids Res. 2017;45:D833–9.

81. Brown AS, Patel CJ. A review of validation strategies for computational drug repositioning. Brief Bioinform. 2018;19:174–7.

82. Paik H, Chen B, Sirota M, Hadley D, Butte AJ. Integrating Clinical Phenotype and Gene Expression Data to Prioritize Novel Drug Uses. CPT Pharmacometrics Syst Pharmacol. 2016;5:599–607.

83. Zhang X, Yuan Z, Ji J, Li H, Xue F. Network or regression-based methods for disease discrimination: a comparison study. BMC Medical Research Methodology. 2016;16:100.

84. Liu H, Song Y, Guan J, Luo L, Zhuang Z. Inferring new indications for approved drugs via random walk on drug-disease heterogenous networks. BMC Bioinformatics. 2016;17:539.

85. Maciukiewicz M, Marshe VS, Hauschild A-C, Foster JA, Rotzinger S, Kennedy JL, et al. GWAS-based machine learning approach to predict duloxetine response in major depressive disorder. Journal of Psychiatric Research. 2018;99:62–8.

86. Zitnik M, Zupan B. Jumping across biomedical contexts using compressive data fusion. Bioinformatics. 2016;32:i90–100.

87. Carson MB, Lu H. Network-based prediction and knowledge mining of disease genes. BMC Medical Genomics. 2015;8:S9.

88. Gu C, Liao B, Li X, Li K. Network Consistency Projection for Human miRNA-Disease Associations Inference. Sci Rep. 2016;6. doi:10.1038/srep36054.

89. Detector Performance Analysis Using ROC Curves | Receiver Operating Characteristic | Signal To Noise Ratio. Scribd. https://es.scribd.com/document/339719122/Detector-Performance-Analysis-Using-ROC-Curves. Accessed 3 Sep 2018.

90. Bustin SA. The reproducibility of biomedical research: Sleepers awake! Biomolecular Detection and Quantification. 2014;2:35–42.

91. Kobourov SG. Spring Embedders and Force Directed Graph Drawing Algorithms. arXiv:12013011 [cs]. 2012. http://arxiv.org/abs/1201.3011. Accessed 3 Sep 2018.

92. Shannon P, Markiel A, Ozier O, Baliga NS, Wang JT, Ramage D, et al. Cytoscape: A Software Environment for Integrated Models of Biomolecular Interaction Networks. Genome Res. 2003;13:2498–504.

93. Le D-H, Pham V-H. HGPEC: a Cytoscape app for prediction of novel disease-gene and disease-disease associations and evidence collection based on a random walk on heterogeneous network. BMC Systems Biology. 2017;11:61.

94. Piñero J, Queralt-Rosinach N, Bravo À, Deu-Pons J, Bauer-Mehren A, Baron M, et al. DisGeNET: a discovery platform for the dynamical exploration of human diseases and their genes. Database (Oxford). 2015;2015:bav028.

95. Perfetto L, Briganti L, Calderone A, Cerquone Perpetuini A, Iannuccelli M, Langone F, et al. SIGNOR: a database of causal relationships between biological entities. Nucleic Acids Res. 2016;44:D548–54.

96. Lo Surdo P, Calderone A, Iannuccelli M, Licata L, Peluso D, Castagnoli L, et al. DISNOR: a disease network open resource. Nucleic Acids Res. 2018;46:D527–34.

97. Calderone A, Cesareni G, Stegle O. SPV: a JavaScript Signaling Pathway Visualizer. Bioinformatics. 2018;34:2684–6.

98. Pavlopoulos GA, Paez-Espino D, Kyrpides NC, Iliopoulos I. Empirical Comparison of Visualization Tools for Larger-Scale Network Analysis. Advances in Bioinformatics. 2017. doi:10.1155/2017/1278932.

99. Westergaard D, Stærfeldt H-H, Tønsberg C, Jensen LJ, Brunak S. A comprehensive and quantitative comparison of text-mining in 15 million full-text articles versus their corresponding abstracts. PLOS Computational Biology. 2018;14:e1005962.

100. Valle EPG del, García GL, Santamaría LP, Zanin M, Ruiz EM, González AR. Evaluating Wikipedia as a Source of Information for Disease Understanding. In: 2018 IEEE 31st International Symposium on Computer-Based Medical Systems (CBMS). 2018. p. 399–404.

101. Grewal N, Singh S, Chand T. Effect of Aggregation Operators on Network-Based Disease Gene Prioritization: A Case Study on Blood Disorders. IEEE/ACM Transactions on Computational Biology and Bioinformatics. 2017;14:1276–87.

102. Karahalil B. Overview of Systems Biology and Omics Technologies. Current medicinal chemistry. 2016;23.

